# Pathogenic *DVL* frameshifting variants in Robinow syndrome disrupt WNT signaling and cellular dynamics

**DOI:** 10.1101/2025.08.02.668297

**Authors:** Chaofan Zhang, Rituparna Sinha Roy, Ming Yin Lun, Juliana F. Mazzeu, Janson White, Wu-Lin Charng, Nathaniel Peters, Jonas A. Gustafson, Harshini Iyer, Zain Dardas, Chad A. Shaw, Hyun Kyoung Lee, V. Reid Sutton, James R. Lupski, Claudia M.B. Carvalho

## Abstract

Robinow syndrome (RS) is a genetically heterogeneous rare disorder involving six genes in the WNT/planar cell polarity (PCP) signaling pathway. Frameshifting variants in *DVL* genes that introduce a novel basic C-terminus are a common cause of autosomal dominant RS, accounting for ∼33% of individuals without *ROR2* variants. Here, we review ClinVar and literature variants affecting *DVL* paralogs resulting in RS and investigate the cellular effects of pathogenic *DVL* frameshift variants with mutant tails replacing at least 82 amino acids. In silico analysis of the new C-termini revealed altered intrinsically disordered regions (IDRs), charge distribution, and predicted protein structures. To explore potential altered biological mechanisms caused by novel C-termini, we generated wild-type (WT), frameshift, and truncated constructs of *DVL1-3*, and analyzed their behavior in a transfection-based in vitro systems. DVL proteins normally polymerize into cytoplasmic puncta that redistribute upon WNT stimulation. Immunocytochemistry showed that mutant DVL proteins failed to change their localization in response to WNT ligands, in contrast to WT alleles—a consistent observation across all three DVLs. In line with this, TOPFlash reporter assays demonstrated that mutant DVL1 and DVL3 failed to activate canonical WNT signaling, while WT proteins induced strong activation. Additionally, the mutant C-terminal tail interfered with CSNK1E-induced phosphorylation, offering a potential mechanism underlying the impaired WNT response. Our results provide further understanding of the cellular consequences of pathogenic *DVL* frameshifting variants and offers insights into the effects of such alleles on WNT signaling, and the perturbations thereof, that may lead to developmental phenotypes observed in RS.

## Introduction

This study aims to investigate the underlying biological perturbations in WNT signaling due to the presence of frameshifting alleles in *DVL* paralogs, which result in autosomal dominant Robinow syndrome (DRS2, MIM#616331; DRS3, MIM#616894). WNT signaling controls many critical early developmental and post-natal physiological processes in humans and mice (Niehrs 2012; Wiese et al. 2018). Signaling studies identified three main WNT pathways depending upon the receptors: WNT/β-catenin, WNT/Planar Cell Polarity (PCP), and WNT/Calcium (Ca^2+^). WNT/β-catenin is known as the canonical signaling pathway, while the other two are referred to as the non-canonical or WNT/β-catenin-independent signaling pathways.

Both WNT/β-catenin and PCP signaling pathways consist of a highly conserved molecular machinery but have different functions. WNT/β-catenin signaling coordinates cell fate, proliferation, differentiation, and tissue homeostasis (Niehrs 2012; Ross et al. 2005; Wiese et al. 2018) whereas WNT/PCP signaling is essential for vertebrate embryogenesis, including convergent-extension movements, limb bud outgrowth, patterning, and cellular orientation. (Abd-El-Barr et al. 2007; Mlodzik 2002; Ross et al. 2005; Strutt et al. 2011; Tobin et al. 2008; Yang and Mlodzik 2015). The WNT signaling pathway is important during both skeletal development and bone mass maintenance later in life (Huybrechts et al. 2020; Laine et al. 2013; Lojk and Marc 2021). Perturbation of this WNT pathway can cause several skeletal dysplasias (Brandi 2014; Craig et al. 2023; Mortier et al. 2019).

WNT/PCP signaling is crucial during embryonal bone and joint formation. It has been reported that PCP signaling is associated with Wnt5a gradient mediated long bone cartilage elongation along the proximal to distal (P-D) axis (Duan et al. 2022; Li and Dudley 2009; Wang et al. 2011), where it was shown to be involved in osteoblast precursors migration and differentiation in mice (Wan et al. 2018). Pathogenic variants in genes that are crucial for WNT/PCP signaling have been linked to human diseases, such as neural tube defects (MIM#182940) (Allache et al. 2012; De Marco et al. 2014; Lei et al. 2010; Ross et al. 2005; Tobin et al. 2008), epilepsy (MIM#612437) (Bassuk et al. 2008; Tao et al. 2011), and autism (Sowers et al. 2013; Tao et al. 2011). To date, there are only two inherited skeletal dysplasia conditions associated with PCP signaling (Lojk and Marc 2021; Mortier et al. 2019): the *ROR2* related autosomal dominant brachydactyly type B1 (MIM#113000) (Afzal and Jeffery 2003; Schwarzer et al. 2009), and RS (Person et al. 2010; van Bokhoven et al. 2000; White et al. 2015; White et al. 2018; White et al. 2016; Zhang et al. 2022).

RS is a genetically heterogeneous disorder characterized by skeletal dysplasia and a distinctive facial gestalt and physical appearance, i.e., craniofacial dysmorphology rendering a recognizable pattern of human malformation. Six genes that converge on the WNT/PCP signaling pathway have been associated with RS (*DVL1*, *DVL3*, *FZD2*, *NXN*, *ROR2*, and *WNT5A*) (White et al. 2018). WNT5A acts as a soluble extracellular ligand of ROR2 and FZD2, together they can trigger the DVL orthologs to activate a branch of the non-canonical WNT signaling pathway (Oishi et al. 2003; Sokol 2000; Yang and Mlodzik 2015). *DVL2* has recently been proposed as another candidate RS-associated gene, with two heterozygous likely pathogenic variants reported (Velasco and Cummings 2022; Zhang et al. 2022).

Reflecting the complexity of WNT signaling, the regulation of developmental processes depends on a fine balance between the canonical (β-catenin–dependent) and non-canonical PCP pathways. DVL proteins play a central role in maintaining this equilibrium, where upregulation of one branch typically suppresses the other. For example, mice exhibiting elevated canonical WNT signaling display phenotypes indicative of disrupted PCP signaling such as increased bone density and altered tibial morphology (Gao et al. 2011; Kuss et al. 2014; Lu et al. 2013). Maintaining canonical and non-canonical WNT signaling balance is critical for proper neuronal differentiation and function (Niehrs 2012; Yang 2012), as well as normal bone development and destruction (Lojk and Marc 2021). Consequently, the pathogenesis of RS is thought to result not from a simple loss of WNT/PCP pathway but from a context-dependent perturbation of their dynamic signaling balance.

The dishevelled protein (Dsh) was first discovered in *Drosophila* as a segment polarity gene (*dsh*); three orthologs (*DVL1*, *DVL2* and *DVL3*) were identified in mammals, including humans and mice (Semenov and Snyder 1997). Heterozygous insertions and deletions (indels) in all three orthologs of *DVL* are the most frequent cause of sporadic and autosomal dominant RS (AD-RS) (63.8% in our in-house cohort), and accounts for an estimated 33% of individuals with a clinical diagnosis of RS without pathogenic variants affecting *ROR2* (White et al. 2018; Zhang et al. 2022). Bunn et al. first demonstrated that C-terminal frameshift mutations in *DVL1* result in an osteosclerotic form of RS later confirmed by Shayota et al., linking altered WNT signaling to bone phenotypes (Bunn et al. 2015; Shayota et al. 2020), especially relevant for cranial osteosclerosis. More recently, another study provided functional evidence from *Drosophila* and *chicken* models showing that *DVL1* frameshift mutations disrupt planar cell polarity and skeletal patterning, further supporting a conserved pathogenic mechanism across species (Gignac et al. 2023).

Interestingly, RS disease-associated variant alleles were found in the *DVL* genes that cluster within the penultimate and last exons of each gene, generating a -1 frameshift (*DVL1, DVL2* and *DVL3*) or a +1 frameshift (*DVL2*) RNA transcripts (Supplemental Fig. 1). Such mutant mRNA types are predicted to escape the nonsense-mediated mRNA decay (NMD) surveillance system and generate truncated (DVL1, DVL2 and DVL3) or an elongated (DVL2) protein with a novel C-terminal tail (Coban-Akdemir et al. 2018; White et al. 2015; White et al. 2018; White et al. 2016; Zhang et al. 2022). The new C-termini do not affect the DEP domain, but may disrupt phosphorylation sites that specify DVL functions (Bernatik et al. 2014; Ho et al. 2012), and are necessary to establish and stabilize protein-protein interactions with ROR2 as well as FZD2 (Punchihewa et al. 2009; Tauriello et al. 2012; Witte et al. 2010). Moreover, the C-terminal portion of DVL3 has been previously reported to inhibit canonical Wnt signaling (Bernatik et al. 2014; Witte et al. 2010). Phosphorylation of this region by casein kinase 1 epsilon (CSNK1E) was shown to be essential for the characteristic electrophoretic mobility shift of DVL3. Mutation of these C-terminal phosphorylation sites impaired CSNK1E-induced changes in DVL3 subcellular localization, suggesting that this region plays a regulatory role downstream of CSNK1E (Bernatik et al. 2014). In fact, screw-tail dog breeds show distinct morphological traits overlapping with human clinical phenotypes in RS, have been found to be homozygous for a 1 bp deletion in the last exon of *Dvl2* (Mansour et al. 2018). Analysis of the canine DVL2 variant protein showed that its ability to undergo WNT induced phosphorylation is reduced, suggesting that altered WNT signaling may contribute to the RS-like syndrome in the screw tail breeds (Mansour et al. 2018). A more recent study genotyped 1954 dogs from 15 breeds for the *DVL2* variant, suggesting the *DVL2* variant contributes to a Robinow-like syndrome in dogs, affecting tail, skull, and possibly heart development (Niskanen et al. 2021). Collectively, frameshifting variants producing a novel C-terminus are observed in all three *DVL* gene paralogs in mammals, suggesting this specific kind of allele act in a gain-of-function (GoF) manner rather than haploinsufficiency.

We performed *in vitro* experiments to investigate how the heterozygous pathogenic DVL variants observed in probands with RS affect relevant C-terminal function in cells. Here we studied the effect of those variants in responding to WNT canonical signaling, change in subcellular localization and how that affects phosphorylation induced by CSNK1E. We hypothesized that RS-associated frameshifting alleles generate a novel C-terminal tail that disrupts critical regulatory phosphorylation sites. The observed alterations in subcellular localization and phosphorylation suggest impaired regulation of DVL activity within WNT signaling pathways. Future studies are required to directly assess downstream transcriptional target gene expression, including potential impacts on bone cell differentiation. We anticipate that elevated canonical WNT signaling and perturbed WNT/PCP pathway resulting from a replaced DVL C-termini may underlie the bone development malformations and other RS-associated clinical phenotypes in probands.

## Results

### Frameshifting variants in RS are predicted to generate C-termini with a minimum of 82 replaced amino acid residues in DVL1 and DVL3

Since our genotype-phenotype RS study published in 2022 (Zhang et al. 2022), five new *DVL1* variants (Hu et al. 2022; Smith et al. 2024), one new *DVL2* variant (Velasco and Cummings 2022), and three new *DVL3* variants have been reported in the literature or submitted to ClinVar (Supplemental Fig. 1). In total, 37 DVL variants are classified as pathogenic or likely pathogenic in DRS; majority of which (34/37, 92%) are located in the penultimate or last exons consistently predicted to result in a -1 frameshift (Supplemental Fig. 1). Exceptions include one *DVL2* variant resulting in +1 frameshift (Zhang et al. 2022), one *DVL3* variant (c.292del) located in exon 3, and a nonsense variant affecting exon 13 (c.1473C>G). Frameshift variants in *DVL1* are predicted to produce a mutant C-terminal tail of at least 106 amino acids, while those in *DVL3* are predicted to produce a mutant tail of at least 82 amino acids (Supplemental Fig. 2).

To investigate the potential pathogenic roles of frameshift variants in DVL genes, we analyzed the structural and sequence features of the WT and mutant proteins. The C-terminal tails of DVL proteins are highly conserved (Supplemental Fig. 2) and are predicted to be part of an intrinsically disordered region (IDR), as revealed by ANCHOR2 and IUPred2 analyses. Pathogenic frameshift variants are predicted to disrupt the IDR in all DVL genes (Fig. 1A1, B1, C1). Additionally, the frameshift variants altered the residue composition and charge distribution of the DVL C-terminal tails (Fig. 1A2, B2, C2). Notably, the novel C-terminal tails of DVL1 and DVL2 were more basic compared to their WT counterparts, with basic amino acid proportions of 19.86% and 13.73%, respectively, compared to 10.98% and 8.82% in the WT tails. While the basic amino acid proportion in DVL3 (∼15%) remained comparable to the WT, the charge distribution was distinctly altered. To further evaluate the structural impact of these variants, the WT and mutant protein structures were predicted using AlphaFold 2 Colab (Fig. 1A3, B3, C3). All mutant proteins exhibited a loss of the small alpha helix (aa677-685 in DVL1, aa712-724 in DVL2, aa696-704 in DVL3) at the C-terminus observed in the WT proteins, resulting in a looser and disrupted C-terminal structure. The mutant DVL3 also lost a small beta sheet (aa713-715 in DVL3) at more C terminal of WT DVL3 (Fig. 1C3). These structural changes are consistent with the findings from the IDR and charge distribution analyses. Prediction confidence metrics (pLDDT and PAE) are shown in Supplementary Fig. 3.

**Fig. 1.**
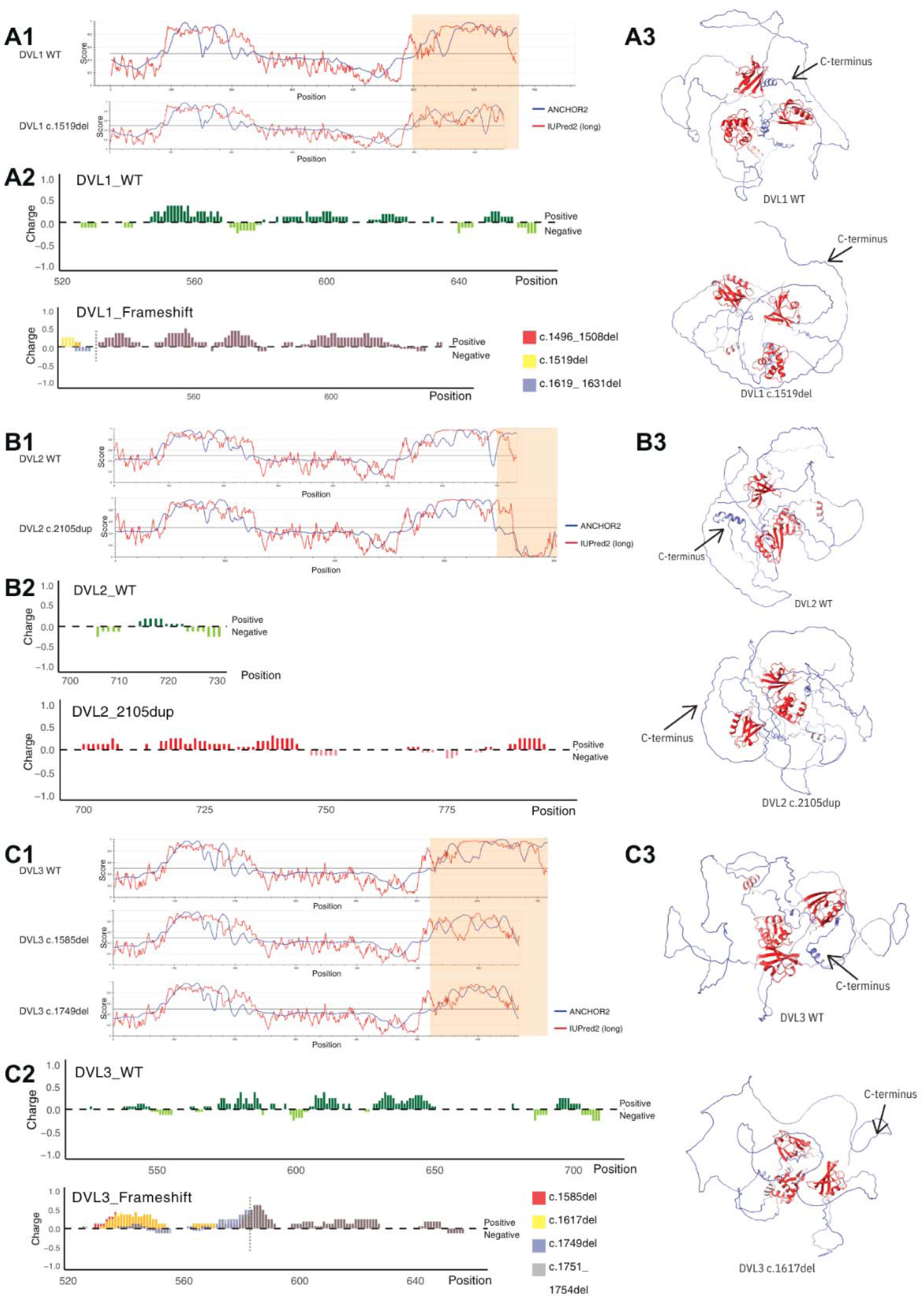
*In silico* prediction of the effects of frameshift variants in DVL genes. The intrinsic disorder region (IDR) prediction, charge distribution analysis, and protein structure modeling were performed for WT and mutant forms of DVL1 (A), DVL2 (B), and DVL3 (C). (A1, B1, C1): Graphs showing the disorder tendency for each residue in WT and mutant DVL proteins. Higher values indicate a greater probability of intrinsic disorder. Disordered regions were predicted using IUPred2A, while disordered binding regions were identified using ANCHOR2. The C-terminal region affected by the frameshift variant is highlighted with an orange box. (A2, B2, C2): Charge plots for the C-terminal regions of WT and mutant DVL proteins as predicted by IDR analysis. Positively charged residues are above zero dashed line, negatively charged residues are below the line, and the boundary of the frameshift variants causing the common mutant C-terminal tail is marked by a gray dashed line. (A3, B3, C3): Structural models of mutant DVL proteins predicted using AlphaFold2 Colab. Structure of WT DVL proteins were obtained from AlphaFold Protein Structure Database (DVL1: AF-O14640-F1-v4; DVL2: AF-O14641-F1-v4; DVL3: AF-Q92997-F1-v4). Arrows indicate the location of the C-terminus.

The C-terminus of DVL1 WT and DVL3 WT displayed similar distributions of small and hydrophobic residues (red), acidic residues (blue), basic residues (magenta), and hydroxyl, sulfhydryl, and amine residues (green) (Supplemental Fig. 2). In contrast, the frameshift variants in the DVL1 and DVL3 mutant tails resulted in an increase in small and hydrophobic as well as basic residues, and a decrease in acidic residues. Additionally, hydroxyl, sulfhydryl, and amine residues were reduced in the DVL1 mutant C-terminus but remained similar ratio in the DVL3 mutant C-terminus. These alterations may disrupt the IDR and the small structural elements at the C-terminus of DVL1 and DVL3.

### *DVL* frameshifting variants in RS guided selection of variants for functional *in vitro* experiments

EBV-transformed Lymphoblastoid cell lines were obtained from patients carrying DVL variants published previously (White et al. 2018): BAB9126 (c.1612_1616dup; p.Ser539Argfs112 variant in *DVL1*, NM_004421.2), BAB9128 (c.1612_1616dup; p.Ser539Argfs112 in *DVL1*), and BAB9236 (c.1608_1623del; p.Ser537Valfs107 in *DVL1*). BAB9127 does not carry *DVL* variants and was used as a negative control. Western blot analysis of these cell lines revealed no detectable DVL protein expression using two different lysis buffers. The anti-DVL antibody was used which could detect all three DVL proteins, anti-GAPDH was used as the positive control (Data not shown).

To investigate the potential biological impact of these C-termini variants in *DVL1* and *DVL3*, we selected one recurrent variant in *DVL1* (c.1519del, DVL1-FS) and two recurrent variants in *DVL3* (c.1749del, DVL3-FS2; c.1585del, DVL3-FS3) identified in RS patients whose phenotypes were validated by our published Human Phenotype Ontology (HPO) based quantitative phenotypic analysis (Fig. 2) (Zhang et al. 2022). To allow comparison with a loss-of-function (LOF) C-terminal allele, truncated versions of DVL1 (c.1519insTAA, DVL1-ST) and DVL3 (c.1749insTAA, DVL3-ST2; c.1585insTAA, DVL3-ST3) lacking the C-terminal tail (losing 164, 133, and 188 amino acids respectively) not observed in probands with RS were also included in the study.

**Fig. 2.**
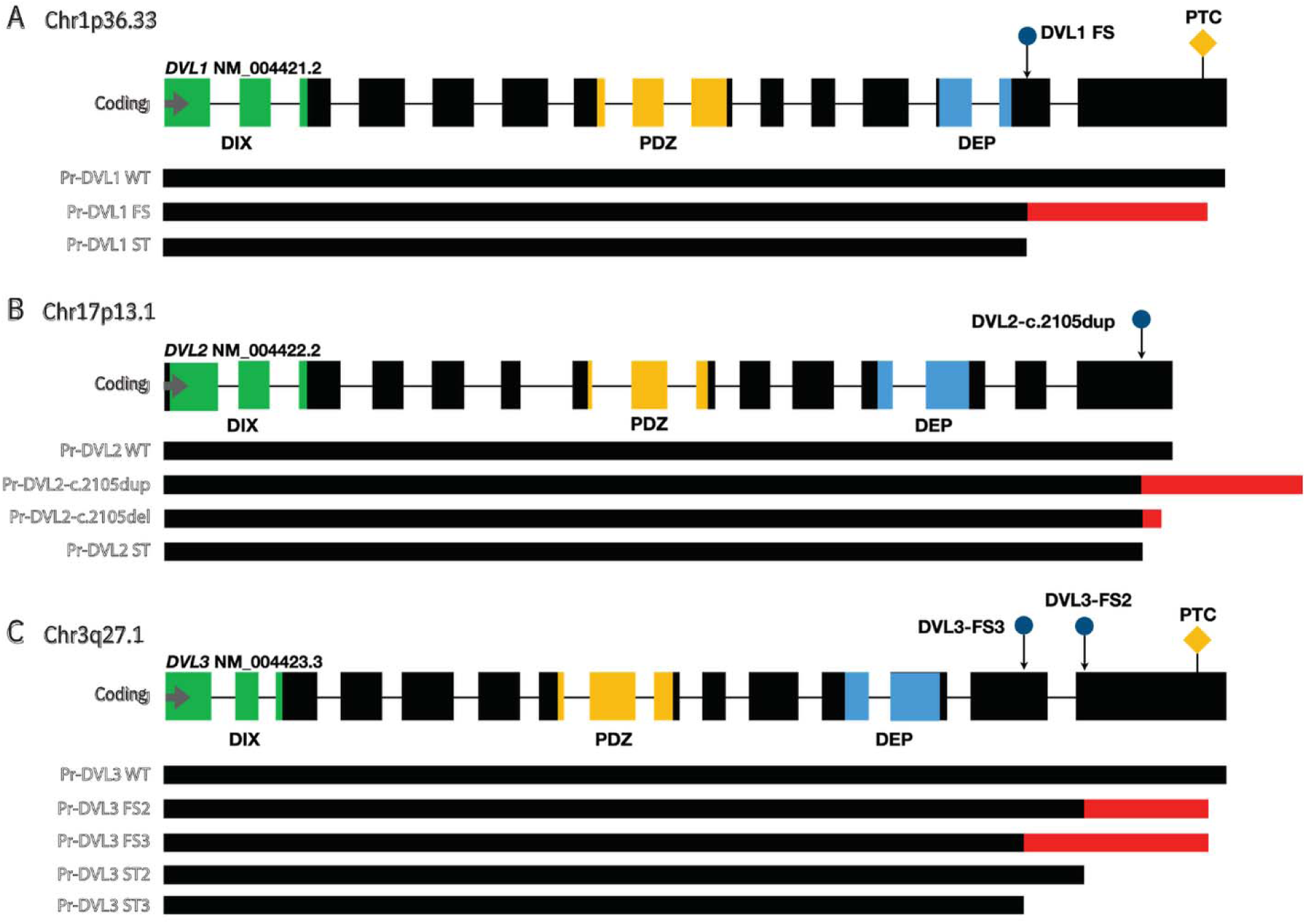
Map location of selected cloned DVL variants and resulting proteins investigated in this study. There are three conserved domains in DVLs that play significant roles in protein/pathway function: a DIshevelled and AXin (DIX) binding domain at the N-terminus; a Post-synaptic density-95, Discs-large and Zonula occludens-1 (PDZ) domain in the mid region of DVL; and a Dishevelled, Egl-10, Pleckstrin (DEP) domain located about midway between the PDZ domain and the C-terminus of DVL. The frameshifting variants identified in RS patients used in this study are shown as blue circles. The chromosome and cytogenetic interval location of the canonical transcripts are provided. The gene structure is depicted with individual exons represented as black rectangles on a horizontal line. The premature termination codon is shown as yellow diamond. Expressed proteins are depicted as dark lines, with black indicating WT regions and red indicating mutant regions of DVL. Three conserved domains are highlighted in different colors: green for the DIX domain, yellow for the PDZ domain, and blue for the DEP domain. (A): The frameshifting variant c.1519delT (DVL1 FS) is located in exon 14 of *DVL1* and is predicted to cause a truncated protein with a novel C-terminal tail. The variant c.1519insTAA (DVL1 ST) is predicted to cause a truncated protein without a C-terminal tail. The C-terminal DEP domain is not predicted to be affected by these variants. (B): The c.2105dup variant (DVL2-c.2105dup) identified in an RS patient is located in exon 15 of DVL2 and is predicted to cause a +1 frameshift, resulting in a longer protein with an additional 102 amino acids in the C-terminal tail. A hypothetical -1 frameshift variant at the same location (DVL2-c.2105del) would result in a shorter protein compared to the canonical one. The variant c.2105insTAA (DVL2 ST) is predicted to cause a truncated protein at the same location without a C-terminal tail. The C-terminal DEP domain is not predicted to be affected by these variants. (C): The frameshifting variants c.1749delG (DVL3 FS2) and c.1585delG (DVL3 FS3) are located in exon 15 and 14 of *DVL3* and are predicted to cause a truncated protein with a novel C-terminal tail. The variant c.1749insTAA (DVL3 ST2) and c.1585insTAA (DVL3 ST3) are predicted to cause a truncated protein without a C-terminal tail. The C-terminal DEP domain is not predicted to be affected by these variants.

To date, only two frameshift variants in *DVL2* have been reported in patients with RS or RS-like phenotypes: one causing a +1 frameshift and the other causing a -1 frameshift (Velasco and Cummings 2022; Zhang et al. 2022). For comparative analysis with *DVL1* and *DVL3*, the +1 frameshift variant in *DVL2* (c.2105dup, DVL2-2105dup) was included, as the phenotype of the proband carrying this variant was validated in our previous HPO phenotypic analysis (Zhang et al. 2022). Additionally, a truncated version (c.2105insTAA, DVL2-ST) and a conceptual -1 frameshift variant (c.2105del, DVL2-2105del) at the same position were generated for further comparison (Fig. 2).

To assess the endogenous expression of DVL proteins, qRT-PCR was performed to analyze the mRNA levels of DVL genes in human HEK293T cells, with normalization to beta-actin (ACTB). The relative expression levels of DVL1, DVL2, and DVL3 were found to be 1.28%, 12.29%, and 1.99% of ACTB, respectively (Supplemental Fig. 4A, 4B). DVL proteins were successfully detected in HEK293T cells overexpressing various DVL constructs (Supplemental Fig. 4C). Furthermore, the effects of WNT ligands were tested in HEK293T cells with or without overexpression of DVL proteins. Expression did not change significantly upon WNT3A and WNT5A under those conditions (Supplemental Fig. 4D-F). Together, these data demonstrate that endogenous DVL protein and RNA levels are low in HEK293T cell, particularly for DVL1 and DVL3, supporting the use of an overexpression system for subsequent functional assays.

### C-terminal integrity of DVL proteins is required for WNT ligands-induced localization shifts

Dvl proteins usually form dynamic multi-protein super-complexes containing Dvl1, Dvl2, and Dvl3 (Lee et al. 2008; Schwarz-Romond et al. 2005), observed in the cell cytoplasm as Dvl puncta (Fig. 3A). A decreased polymerization of Dvl complexes is associated with localization changes, as visualized microscopically as cellular phenotypes from puncta to an even cellular distribution, whereas overexpression of Fzd receptor and WNT signaling stimulation was shown to cause membrane accumulation of Dvl molecules (Axelrod et al. 1998; Bernatik et al. 2014). It has been shown that removal of Dishevelled from the cell membrane disrupts convergent extension of PCP signaling, whereas both membrane and cytoplasm targeted Dishevelled accumulation can activate canonical signaling in vertebrate embryos (Park et al. 2005). Examples of punctate, even distribution and cytosolic accumulation of DVLs observed in our experiments are displayed in Fig. 3A. Full-field images for DVL1, DVL2, and DVL3 are provided in Supplemental Fig. 5.

**Fig. 3.**
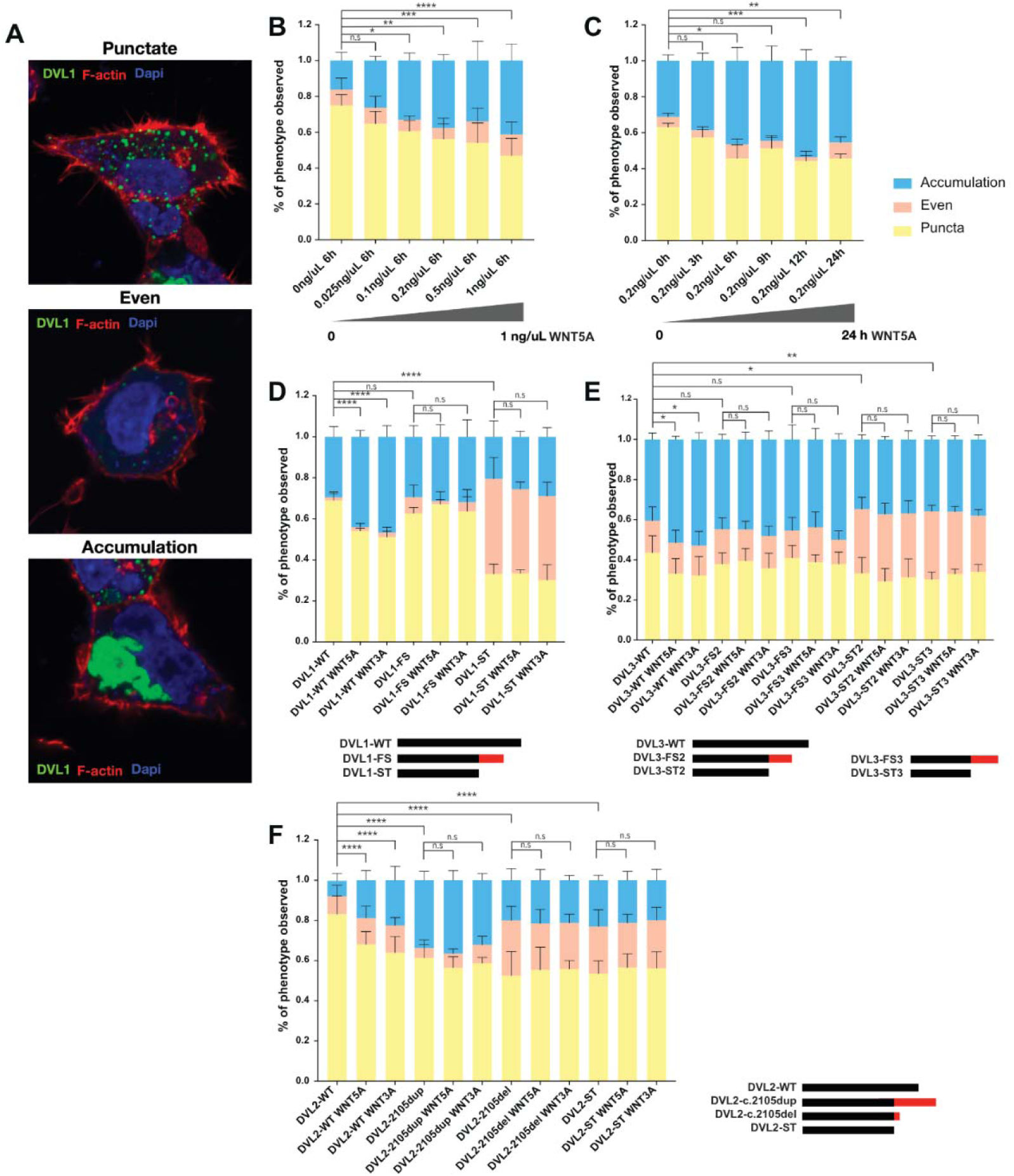
Analysis of subcellular localization of DVL. (A): Three typical distribution patterns of DVL1, punctate, even, and accumulation. Green: GFP-tagged WT DVL1; Red: 650 Phalloidin, label of cytoskeleton; Blue: DAPI, label of nucleus. (B): HEK293T cells were transfected with GFP-tagged DVL1-WT construct and treated with gradient concentration of WNT5A recombinant protein (0-1ng/uL), incubated for 6 hours. The ratio of distribution patterns of DVL1 was calculated and analyzed. Colors indicate different distribution patterns: blue, accumulation; peach, even; yellow, puncta. (C): HEK293T cells were transfected with GFP-tagged DVL1-WT construct and treated with medium containing 0.2 ng/uL of WNT5A recombinant protein, incubated for 0-24 hours. The ratio of distribution patterns of DVL1 was calculated and analyzed. (D): HEK293T cells were transfected with corresponding GFP-tagged *DVL1* constructs, with or without treatment of 0.2 ng/uL of WNT3A or WNT5A recombinant protein, incubated for 6 hours. The ratio of distribution patterns of DVL1 was calculated and analyzed. Black indicates WT coding region of DVL1, red indicates mutant coding region of DVL1. (E): HEK293T cells were transfected with corresponding HA-tagged *DVL3* constructs, DVL3 was stained with anti-HA antibody with or without treatment of 0.2 ng/uL of WNT3A or WNT5A recombinant protein, incubated for 6 hours. The ratio of distribution patterns of DVL3 was calculated and analyzed. Black indicates WT coding region of DVL3, red indicates mutant coding region of DVL3. (F): HEK293T cells were transfected with corresponding Flag-tagged *DVL2* constructs, DVL2 was stained with anti-Flag antibody with or without treatment of 0.2 ng/uL of WNT3A or WNT5A recombinant protein, incubated for 6 hours. The ratio of distribution patterns of DVL2 was calculated and analyzed. Black indicates WT coding region of DVL2, red indicates mutant coding region of DVL2. logistic regression models were conducted for puncta (or even) to compare each condition and WT. Mixed-effects logistic regression (Fig. 3B, 3C) or multi-factor logistic regression (Fig. 3D, 3E, 3F) were fitted including concentration (0-1 ng/uL) and treatment duration (0-24 hours) analyses (Fig. 3B, 3C) as well as genotype (WT, FS, ST), ligand (none, WNT5A, WNT3A), and their interactions (Fig. 3D, 3E, 3F). *P*-values are computed by Wald tests. Post-hoc comparisons were performed against the appropriate reference condition and odds ratios with 95% confidence intervals are determined. For all post-hoc tests, *p*-values were corrected for multiple comparisons using the Dunnett method. Asterisk: significant difference (*p*=0.05) was observed. More than 800 cells were analyzed per condition across two independent experiments (triplicate slides each), ensuring robust quantification of the observed localization patterns. Significance thresholds: *p* < 0.05 (*), *p* < 0.005 (**), *p* < 0.0005 (***), *p* < 0.0001 (****), n.s: no significance.

WNT5A is an extracellular ligand that activates the WNT noncanonical pathway (Maeda et al. 2012). A previous report showed that the binding between Dishevelled and *APC* gene product was enhanced by Wnt5a (Matsumoto et al. 2010). Given that the local concentration and subcellular localization of Dishevelled protein is essential for both canonical and noncanonical pathways (Park et al. 2005), we tested whether WNT3A and WNT5A affect DVL WT and mutant cellular localization.

HEK293T cells were transfected with GFP-tagged DVL1-WT. GFP-tagged *DVL1* constructs originally reported by Bunn et al., 2015, were obtained from Dr. Robertson (Bunn et al. 2015). To further investigate the functional consequences of this variant, we examined its subcellular localization and response to WNT ligand stimulation in comparison with WT *DVL1*. Different concentrations (0-1ng/uL) of WNT5A recombinant protein were added to the media and cells were incubated for 6 hours. We observed predominantly punctate localization (∼74.9%) of DVL1 protein without WNT5A stimulation (Fig. 3B). Increasing concentrations of WNT5A, DVL1 proteins acquired more of a cytoplasmic accumulation pattern and less punctate localization as visualized by fluorescence microscopy. This change is statistically significant (p<0.05, mixed effect logistic model, see Methods) when the WNT5A concentration is higher than 0.1 ng/uL. We next implemented a similar experiment to explore the relationship between DVL1 localization and incubation time. HEK293T cells overexpressed DVL1-WT were incubated in medium containing 0.2 ng/uL WNT5A for 0-24 hours. After more than six hours incubation, DVL1 proteins acquired a cellular phenotype showing significantly more accumulation and less punctate localization (Fig. 3C). Similar experiments were conducted in HEK293T cells overexpressing WT DVL2, WT DVL3, and empty vectors expressing only tags to explore the endogenous settings of DVL localization patterns, with or without stimulation by WNT3A or WNT5A. Similar to WT DVL1, DVL2 and DVL3 can be stimulated to change localization from puncta to accumulation (Supplemental Fig. 6). However, the majority of cells expressing only tags showed even localization, which was not altered by the addition of WNT3A or WNT5A ligands. These results indicate that overexpressed WT DVL proteins are stimulated to cytosolic accumulation by WNT ligands as expected.

To explore whether the frameshifting variants in the C-terminus of *DVL* identified in RS patients influences the DVL subcellular localization, with or without stimulation of the extracellular ligands WNT3A/WNT5A, we used WT GFP tagged *DVL1* construct (DVL1-WT) and GFP tagged *DVL1* construct with the most frequent frameshifting variant found in RS subjects (DVL1-FS, c.1519delT). To compare the results of this allele carrying truncated novel C-terminal tail with an allele that removes the full C-terminal tail, we also included a GFP-tagged *DVL1* construct without the C-terminal tail (DVL1-ST, c.1519insTAA). WT version (DVL3-WT), two different frameshifting versions (DVL3-FS2, c.1749delG; DVL3-FS3, c.1585delG), and two shortened protein versions (DVL3-ST2, c.1749insTAA; DVL3-ST3, c.1585insTAA) of HA-tagged *DVL3* constructs were also generated. For *DVL2* constructs, except the WT version (DVL2-WT), mutant version with the +1 frameshifting variant observed in RS patient (DVL2-2105dup, c.2105dupC), and short version (DVL2-ST, c.2105insTAA), we also included a mutant version with -1 frameshifting variant at the same locus (DVL2-2105del, c.2105delC) to compare. The locations and resulting proteins of these variants are shown in Fig. 2.

Cells were transfected with DVL1-WT, DVL1-FS, and DVL1-ST constructs respectively, DVL1-FS showed a similar localization pattern as DVL1-WT without WNT stimulation: 70% of cells were observed to exhibit puncta for DVL1 proteins whereas ∼30% of the cells showed a cytosolic accumulation pattern. The DVL1-ST showed significantly higher rate of even localization (∼45%) than DVL1-WT and DVL1-FS (Fig. 3D).

A similar series of experiments was conducted using *DVL3* constructs, two different frameshifting versions (FS, RS associated variant alleles) and short versions (ST) were studied. Each of the two DVL3 frameshifting proteins showed a similar localization pattern as DVL3-WT: DVL3 proteins exhibited puncta in ∼39% of cells whereas ∼45% of the cells showed an accumulation pattern. Each of the two short DVL3 showed significant more even localization (∼34%) than DVL3-WT (∼15%) (Fig. 3E).

The results of analogous experiments using *DVL2* constructs are shown in Fig. 3F. Most of the DVL2-WT exhibited puncta (>80%), and ∼8% showed accumulation localization (Fig. 3F). DVL2-2105dup showed significantly more accumulation (∼34%) and less puncta (∼61%) compared to DVL2-WT. Both DVL2-2105del and DVL2-ST showed significantly more even localization (∼24%), similar to DVL1-ST and DVL3-ST.

Quantification was performed based on classification of cells into puncta, even, or accumulation patterns as described in Methods. After adding WNT3A or WNT5A, DVL1-WT showed significantly less puncta (∼53%) and more accumulation (>45%) localization in comparison to cells without WNT accumulation (Fig. 3D). However, after adding WNT3A or WNT5A, DVL1-FS showed a deficit in WNT-stimulated localization change (puncta to accumulation). The specific pattern of DVL1-ST (more even) did not change with WNT stimulation either (Fig. 3D). Importantly, altered subcellular distribution of *DVL1* frameshift variants has also been reported using N-terminally FLAG-tagged constructs in HEK293 cells (Gignac et al. 2023), supporting that these localization differences are intrinsic to the variant proteins rather than specific to a particular tagging strategy.

Similar to DVL1, DVL3-WT showed significantly less puncta (∼32%) and more accumulation (>51%) localization after WNT stimulation. However, each of the two DVL3 frameshifting proteins showed a deficit in WNT-stimulated localization change (puncta to accumulation). The even pattern of DVL3-ST was not changed after WNT stimulation (Fig. 3E).

After WNT stimulation, DVL2-WT showed significantly less puncta (∼66%) and more accumulation (>20%) localization (Fig. 3F). No further localization changes of DVL2-2105dup, DVL2-2105del, or DVL2-ST were observed after WNT stimulation.

In aggregate, these results suggests that the human pathogenic alleles leading to frameshifting in the DVL protein are not responsive to WNT3A and WNT5A stimulus *in vitro*. Moreover, these data support the hypothesis that the C-terminal tail of DVL is essential for the localization of the DVL supramolecular complex.

### DVLs with a novel C-terminus are insensitive to WNT stimulus when co-overexpressed with DVL WT

Our previous results indicate that the pathogenic DVL C-terminal tail interferes with the dynamic location changes expected upon WNT stimulation, i.e. less puncta and more accumulation. We hypothesize that mislocalization of the DVL super complex disrupts WNT signaling transduction within the cells. To investigate the effect of the *DVL* variants on the activity of the WNT canonical signaling pathway, we quantified canonical WNT signaling utilized a transient transfection system using the TOPFlash reporter assay (Kumar et al. 2008). Non-canonical PCP signaling lacks a consensus transcriptional reporter; accordingly, we assessed PCP-responsive behavior through ligand-induced DVL relocalization rather than a luciferase readout. Luciferase assays were analyzed separately for each DVL paralog (DVL1, DVL2, DVL3), comparing WT, frameshift, and truncated constructs under identical conditions (Fig. 4).

**Fig. 4.**
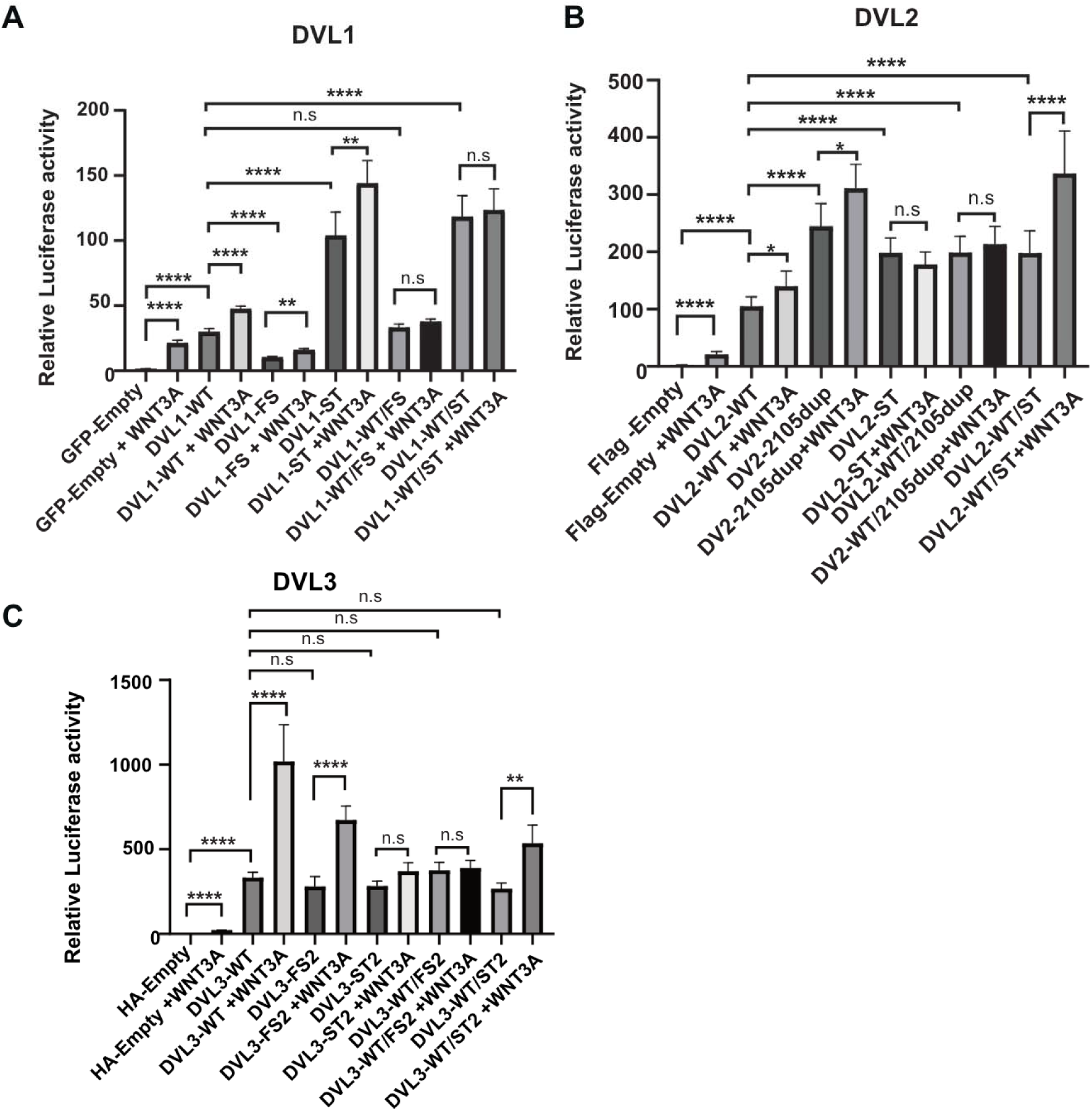
Impact of *DVL* constructs on WNT canonical signaling. (A–C): HEK293T cells were transientl transfected with SuperTopFlash and FOPFlash reporters together with a fixed amount of the indicated *DVL* constructs or with a 1:1 stoichiometric ratio of two constructs while keeping the total amount of DVL plasmid constant. (A): GFP-tagged *DVL1* constructs. (B): FLAG-tagged *DVL2* constructs. (C): HA-tagged *DVL3* constructs. For all panels, each bar represents the mean of three independent biological replicates, with each biological replicate consisting of a separate transfection performed in triplicate wells. Error bars indicate the standard error of the mean. Relative STF/FOP values was analyzed using a linear mixed-effects model with log2-transformed activity as the outcome variable, genotype, ligand, and their interaction as fixed effects, and experiment as a random effect. P-values are computed by Wald tests. Significance thresholds: *p* < 0.05 (*), *p* < 0.005 (**), *p* < 0.0005 (***), *p* < 0.0001 (****), n.s: no significance.

The firefly reporter plasmids for SuperTopFlash together with a fixed amount of each *DVL1* construct or with a 1:1 stoichiometric ratio of two *DVL1* constructs were transfected into HEK293T cells, with or without stimulation of WNT3A. The effects of DVL variants on canonical WNT signaling were assessed by measuring luminescence activity. Cells transfected with the reporter and GFP tagged empty construct were used as a negative control. As shown in Fig. 4A, transfection of DVL1-WT led to an increase of WNT canonical activity, and adding WNT3A further elevates it (∼59%) significantly. Transfection of DVL1-FS also elevated canonical activity, but to a significantly lesser extent than DVL1-WT. In contrast, DVL1-ST significantly increased canonical activity more than DVL1-WT. Further increase in activity was observed when the ligand WNT3A was added.

To test the stoichiometric balance between WT and mutant allele, we co-transfected DVL1-WT and DVL1-FS in a 1:1 ratio while maintaining the same total amount of the construct as each alone. A similar 1:1 ratio of DVL1-WT and DVL1-ST transfection was also examined. DVL1-WT:DVL1-FS combination showed comparable level as DVL1-WT alone, whereas a significantly higher elevated canonical activity was found in DVL1-WT:DVL1-ST combination in comparison to DVL1-WT alone. However, no further elevation was observed in the presence of WNT3A for both combinations (Fig. 4A).

Similar experiments were conducted by using HA-tagged *DVL3* constructs, a frameshifting *DVL3* construct, DVL3-FS2 and DVL3-ST, were tested. As shown in Fig. 4C, transfection of DVL3-WT also led to an increase of WNT canonical activity, and significantly higher canonical activity was observed with the WNT stimulus WNT3A (almost 3 times). Transfection of DVL3-FS2 or DVL3-ST showed a similar canonical activity as DVL1-WT, and significantly more elevation was observed in DVL3-FS2 when WNT3A added. However, no further elevation in activity was observed in DVL3-ST when adding WNT3A. A similar elevated canonical activity was found in DVL3-WT:DVL3-FS2 and DVL3-WT:DVL3-ST combinations in comparison to DVL3-WT alone, and significantly higher elevated canonical activity was found in DVL3-WT:DVL3-ST when WNT3A was added. Similar to that observed with the DVL1-FS, no further elevation was observed in the presence of WNT3A in DVL3-WT:DVL3-FS2 (Fig. 4C).

The Flag-tagged *DVL2* constructs, including +1 frameshifting (DVL2-2105dup) and DVL2-ST, were also tested. As shown in Fig. 4B, transfection of DVL2-WT led to an increase of WNT canonical activity, and higher canonical activity was observed with the WNT stimulus WNT3A. However, all other conditions tested showed a higher level than DVL2-WT, and DVL2-WT:DVL2-2105dup is insensitive to WNT3A.

In summary, both DVL frameshift variants and DVL short variants show canonical signaling activation on their own or when co-overexpressed with DVL WT. However, the phenotypes differ between frameshift and short variants. In all, frameshift variants in *DVL1*, *DVL2* and *DVL3* alone exhibit increased canonical activity when stimulated by WNT3A, while a combination of frameshift variants and WT do not change luciferase activity. Importantly, unlike DVL WT, DVL frameshift variants are generally insensitive to WNT stimulus when co-overexpressed with DVL WT.

### Variants in the C-terminus of DVL interfere with CSNK1E induced electrophoretic migration of DVL

We hypothesized that frameshifting variants in *DVL* genes result in generation of a C-terminus without some of the phosphorylation sites essential for electrophoretic mobility of DVL The latter were shown to be relevant for subcellular localization and function (Ho et al. 2012). To test this hypothesis, we transfected casein kinase 1 Epsilon (CSNK1E), the major kinase responsible for WNT-induced Dvl3 phosphorylation (Bernatik et al. 2014) to 293T cells overexpressed DVL proteins. CSNK1E was shown to be responsible for phosphorylation of some sites of the C-terminal tail of Dvl3 that are conserved in all Dvl (amino acid at position 567, p.567), Dvl1 (p.611-612), or Dvl2 (p.689), which were affected in most of the frameshifting variants observed in RS patients (Bernatik et al. 2014).

Introduction of negative charges by phosphorylation decreases the binding of SDS (sodium dodecyl sulfate) to phospho-proteins and retards the electrophoretic migration. To test if the frameshifting variants in *DVL* remove phosphorylation sites and change electrophoretic migration, HEK293T cells were transfected with WT, frameshifting, and short versions of *DVL* constructs, or co-transfected with Flag-tagged CSNK1E construct together with the *DVL* constructs, after a 24-hour incubation, whole protein was extracted and used for Western blots. Cells transfected with *DVL* constructs and Flag-tagged empty construct were used as the negative control. Lambda phosphatase (LP) treatment was used to confirm that changes in migration were due to phosphorylation rather than other post-translational modifications.

We observed a shift in the electrophoretic mobility of DVL1-WT and DVL3-WT when co-transfected with CSNK1E, whereas no shift occurred when an empty vector was co-transfected (Fig. 5, Supplemental Fig. 7). This mobility shift, indicative of a post-translational modification, was abolished by LP treatment, confirming that it was CSNK1E-dependent phosphorylation. In contrast, frameshift versions of DVL1 and DVL3 did not exhibit a CSNK1E-induced mobility shift, supporting the interpretation that critical phosphorylation sites are lost in the truncated proteins.

**Fig. 5.**
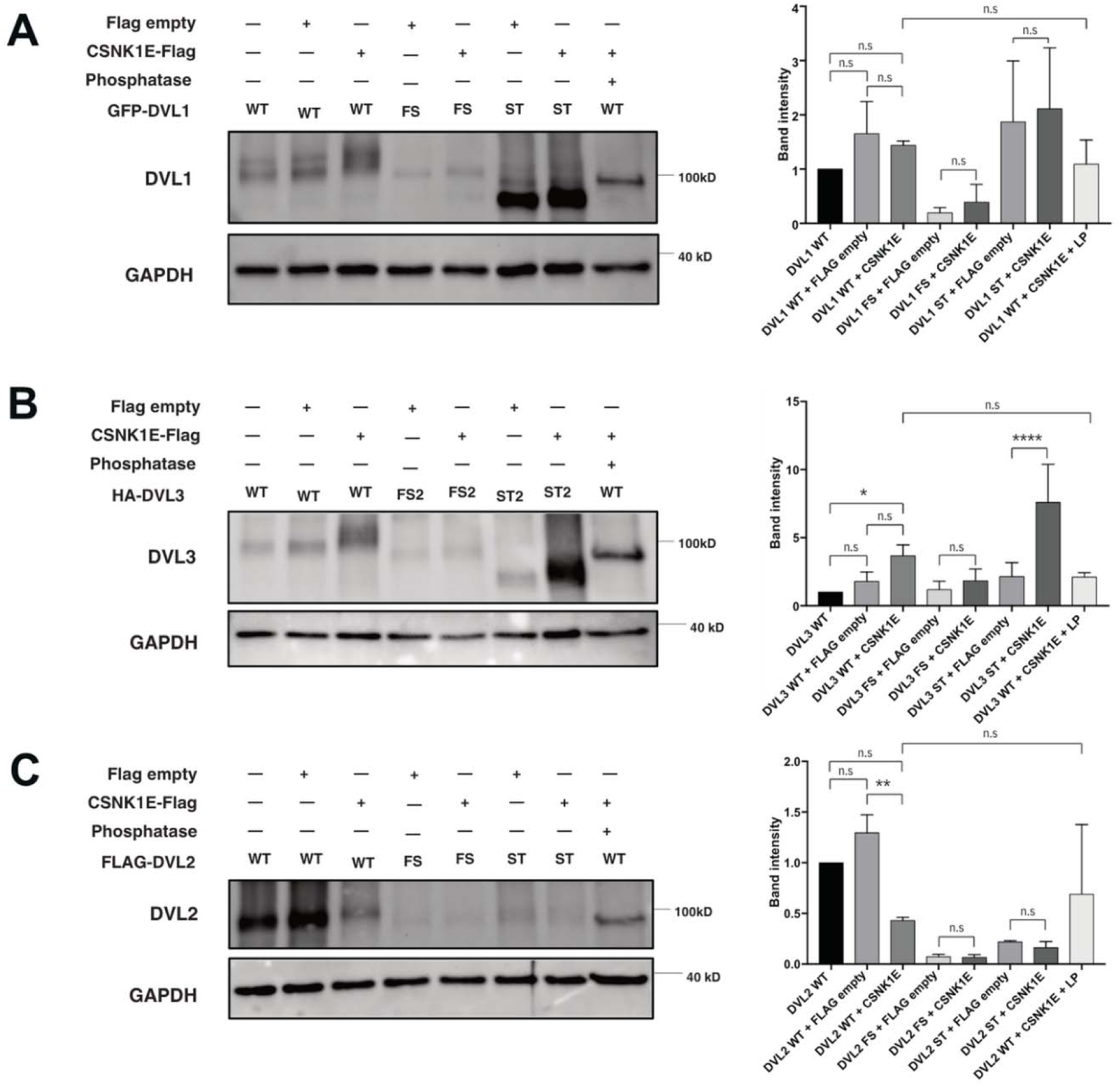
CSNK1E induced electrophoretic migration and phosphorylation dependent shift of DVL. DVL proteins were analyzed by Western blot, and band intensities were quantified using Fiji (ImageJ). For each DVL isoform, three independent biological replicates were performed. Densitometry values were first normalized to the loading control and then expressed relative to the WT condition (set to 1). Data are presented as mean ± SEM. Statistical significance was assessed using unpaired two-tailed Student’s *t*-tests in GraphPad Prism 7. (A): Wester blot result for whole proteins extracted from HEK293T cells transfected with GFP-tagged DVL1-WT, DVL1-FS, DVL1-ST alone, or co-transfected with Flag-tagged CSNK1E or Flag empty construct with the *DVL1* constructs. (B): Western blot result for whole proteins extracted from HEK293T cells transfected with HA-tagged DVL3-WT, DVL3-FS2, DVL3-FS3, DVL3-ST alone, or co-transfected with Flag-tagged CSNK1E or Flag empty construct with the *DVL3* constructs. (C): Western blot result for whole proteins extracted from HEK293T cells transfected with FLAG-tagged DVL2-WT, DVL2-2105dup, DVL2-ST alone, or co-transfected with Flag-tagged CSNK1E or Flag empty construct with the *DVL2* constructs. Beta actin is served as internal control. Lambda phosphatase was added to confirm the migration is caused by phosphorylation. Relative band intensity was analyzed using a linear mixed-effects model, genotype, CSNK1E, and their interaction as fixed effects, and experiment as a random effect. Significance thresholds: *p* < 0.05 (*), *p* < 0.005 (**), *p* < 0.0005 (***), *p* < 0.0001 (****), n.s: no significance.

Quantification of band intensities further revealed that DVL3-WT displayed significantly increased protein abundance when co-transfected with CSNK1E compared with DVL3-WT, suggesting that phosphorylated DVL3 is more stable (Fig. 5B). A similar increase was observed for DVL3-ST when co-transfected with CSNK1E. However, the expression levels of the DVL3 frameshift variants remained consistently low and were unaffected by CSNK1E co-expression.

For DVL2-WT, we detected only a modest mobility shift upon CSNK1E co-transfection, accompanied by a reduction in overall protein abundance. LP treatment abolished the shift, supporting phosphorylation dependence. Both DVL2-FS and DVL2-ST exhibited consistently low expression levels and failed to show a CSNK1E-induced mobility shift. The stability of phosphorylated DVL2 differed markedly from that of DVL3, suggesting paralog-specific effects of phosphorylation. Further investigation is needed to clarify this observation.

Together, these findings indicate that the RS-associated frameshift variants in *DVL1*, *DVL2*, and *DVL3* disrupt CSNK1E-mediated phosphorylation, potentially providing a mechanistic explanation for their pathogenic effects.

## Discussion

To date, 22 *DVL1* variants, 2 *DVL2* variants, and 11 *DVL3* variants associated with an RS phenotype have been identified that follow a specific pattern: they are located in the last or penultimate exon, predicted to cause transcripts that escapes NMD, and result in mutant proteins with a novel C-terminal tail. Of these 35 identified *DVL* variants, 83% (29/35) are indels and 17% (6/35) are splice-site variants utilizing acceptor sites downstream of the canonical site. The novel amino acid residues are found to alter the charge composition, distribution of different types of amino acids, and overall structure of the C-terminus, specifically disrupting the C-terminal IDR (Fig. 1). Similar to our findings, a previous study cataloged over 200,000 genetic variants in C-terminal IDRs and highlighted how frameshift-induced charge alterations can disrupt protein dynamics and localization, contributing to disease mechanisms (Mensah et al. 2023). Although the exact downstream consequences may differ, both studies underscore the critical role of IDRs in maintaining protein functionality and their vulnerability to frameshift variants. A previous study proposed a model in which a disulfide bond between the PDZ domain and the C-terminus contributes to DVL autoinhibition (Qi et al. 2017). We used a similar structural prediction approach with AlphaFold models to visualize the C-terminal changes in the frameshift variants. However, AlphaFold models do not include disulfide bond information, and the C-terminal regions showed low pLDDT confidence, suggesting high disorder, which limits the ability to directly assess whether such autoinhibitory interactions are preserved in the frameshift variants. Because AlphaFold relies heavily on evolutionary information and homologous sequence alignments, structural modeling of these novel C-terminal tails is inherently limited. Therefore, our current data are insufficient to predict specific disulfide bond formation, and our hypothesis instead focuses on C-terminal alteration and abnormal phosphorylation rather than PDZ or DEP domain disruption. Full AlphaFold2 prediction confidence metrics are provided in Supplementary Fig. 3.

Using an *in vitro* cellular transfection system and measuring the observed cellular phenotypes, we provide compelling evidence that the Robinow syndrome associated frameshifting variants in *DVL* interfere with the expected intracellular localization change of the DVL super complex under WNT stimulus. Consistently, the pathogenic mutant alleles in combination with WT do not cause elevated luciferase activity in HEK293T cells in the absence of WNT ligands stimulation supporting disruption of the WNT canonical activity as well. Moreover, the electrophoretic mobility shifts induced by CSNK1E phosphorylation are absent in those mutants.

Overall, these functional studies suggest that DVL FS mutants are unresponsive to WNT3A or WNT5A stimuli, consistent with the functional effects of biallelic LOF variants in ROR2 *in vitro* (Lima et al. 2022). This resemblance extends to recent findings in FZD2 variants, which impair Fzd2–Dvl1 interaction and WNT signal propagation (Punchihewa et al. 2009; Tauriello et al. 2012; Zhang et al. 2022). Our finding provides clues to the cellular effects produced by pathogenic frameshifting DVL variants in RS, supporting a hypothesis of decreased capacity of WNT ligands to transmit the signaling either due to the hypofunction of the ligand (WNT5A) (Hosseini-Farahabadi et al. 2017), lack of the membrane receptor (ROR2) (Lima et al. 2022), truncated or mutated co-receptor that does not pass on the signal (FZD2) (Tauriello et al. 2012), or the mediators that do not localize correctly after WNT stimulation (DVL).

The pathogenicity of frameshifting variants in *DVL* may be partially explained by removal of the C-terminal phosphorylation sites. The C-terminus of DVL has the conserved DEP domain that is activated in PCP signaling, which is required for membrane recruitment via WNT mediated signaling (Pan et al. 2004). Though these frameshifting variants seem to not affect the DEP domain (Fig. 2), the C-terminus carries relevant phosphorylation sites that specify DVL functions (Bernatik et al. 2014; Ho et al. 2012), and are necessary to establish and to stabilize protein-protein interactions with ROR2 as well as FZD2 (Punchihewa et al. 2009; Tauriello et al. 2012; Witte et al. 2010). It has been reported that the hyperphosphorylation of DVL3 requires C-terminal serine clusters, otherwise, the absence of clusters alters the DVL3 subcellular localization and DVL polymerization. Further evidence in support of this contention is the interaction between phosphorylated DVL3 and the non-canonical WNT receptor ROR2 (Nishita et al. 2010). Additionally, the DVL3-ROR2 interaction depends on the DVL C-terminus (Witte et al. 2010), which is lost in all observed *DVL* mutants we have identified thus far (Zhang et al. 2022). Pathogenic variants of phosphorylation sites in the C-terminus of Dvl3 were found to prevent the CSNK1E induced electrophoretic mobility shift and subcellular localization of Dvl3 (Bernatik et al. 2014). It is also possible that the C-terminal phosphorylation can affect the intramolecular interaction between the PDZ domain and DVL C-terminus, which is further supported by a previous report (Śmietana et al. 2011). Both mutant versions of DVL, i.e. FS with a novel C-terminus and ST lacking the C-terminus show similar *in vitro* phenotype of lack of response to WNT stimulus.

However, ST shows a consistent distinct mislocalization phenotype compared to FS. As shown in Fig. 3, the frameshifting DVL proteins exhibited puncta and accumulation localization that is not different from the WT without WNT stimulus. However, the short DVL proteins shows significantly more even localization, perhaps reflecting a less stable protein. Therefore, lack of phosphorylation may not be sufficient to explain the lack of WNT transduction cellular phenotype. Moreover, the fact that all 21 *DVL1* and 8 *DVL3* variants identified in probands with validated RS phenotypes (Zhang et al. 2022) have a potentially similar effect, i.e. producing a new truncated C-terminus, support the hypothesis that full deletion of the C-terminal may not lead to the same disease. This distinct mislocalization phenotype of FS variants is consistently observed across all three DVL paralogs.

It was previously reported that cytoplasmic DVL2 puncta observed under strong overexpression likely represent protein aggregates and are not required for canonical WNT activation (Smalley et al. 2005). In our study, DVL localization was examined under moderate expression levels, focusing on RS–associated frameshift variants. The punctate, even, and accumulated patterns reflect localization states of DVL that dynamically respond to WNT stimulation. The punctate pattern in WT DVLs is reversible after WNT3A or WNT5A treatment, indicating that it represents functional assemblies rather than non-specific aggregates. In contrast, the RS-associated variants fail to redistribute properly, suggesting that C-terminal disruptions impair normal DVL complex dynamics.

CSNK1E co-transfection caused mobility shifts in DVL1-WT and DVL3-WT, but it is not clear in DVL2-WT, suggesting a paralog-specific regulation. The casein kinase 1 σ and e were shown to be essential for both WNT canonical pathway activation and Dvl phosphorylation (Bryja et al. 2007). It is possible that the CSNK1E is not the major kinase for DVL2, and the tests of CSNK1σ and other kinases can potentially provide further insights into underlying mechanisms of the WNT signaling activation and DVL phosphorylation. Until now, only two RS patients with variants in *DVL2* were reported, additional investigation of RS individuals carrying *DVL2* variants is necessary to confirm our findings.

Typically, WT DVL oligomerization to form puncta in cells is a necessary property for canonical WNT signaling (Schwarz-Romond et al. 2005). A study conducted by Bernatik *et al*. indicated that Dvl3 showed predominantly a punctate localization pattern in HEK293T cells, whereas mutant for several important phosphorylation sites all showed a similar localization pattern to that observed with Dvl3 WT (Bernatik et al. 2014). Consistently, our data showed that the mutant DVL1/DVL3 was able to form DVL complex, but the ST did not. Our results also indicate that the +1 frameshifting mutant of DVL2 showed significantly more accumulation and less puncta localization, which is distinguishable from the -1 frameshifting mutant DVL2, the localization of the latter is similar to that which was observed with short DVL2. This can be explained by the consequence of the variants, the +1 variant that was found in the RS individual may generate a longer mutant tail than WT protein, whereas the -1 variant is predicted to cause a much shorter mutant tail, which is similar to the shorter protein (ST) (Fig. 2B). For DVL2, we cannot rule out the contribution of the endogenous protein as a confounder factor.

Co-expression of mutant and WT DVL1 resulted in elevated canonical WNT signaling, consistent with previous findings (Bernatik et al. 2014; Bunn et al. 2015). Bunn et al. demonstrated that *DVL1* frameshift variants exhibit reduced canonical activity when expressed alone but enhance WNT signaling when co-expressed with WT DVL1. Similarly, we observed that mutant constructs alone show diminished activation relative to WT. In our experiments co-expression of FS and WT DVL1 does not result in significantly more elevated canonical WNT signaling, but rather lack of luciferase activation upon WNT3A stimulus. The difference between our results may be due to different transfections systems.

The present study extends these findings in several important ways. First, we studied WT, frameshift and truncated alleles with and without WNT ligands which allowed us to investigate dynamic changes in localization and luciferase assay to signaling responses. The results support our hypothesis that the novel C-terminal tail has a functional role in WNT signaling perturbation in RS. Second, we assessed responses under WNT3A or WNT5A stimulation, further supporting altered regulatory dynamics in both canonical and noncanonical readouts. Third, in addition to *DVL1* alleles we examined *DVL2* and *DVL3* pathogenic variants, highlighting common responses but also a few paralog-specific differences in signaling behavior.

Overexpression of DVLs has been shown to mimic WNT stimulation (Lee et al. 2008), but mutants failed to respond further to WNT, highlighting an impaired feedback mechanism. ST mutants were unable to form DVL complexes or respond to WNT, whereas FS mutants formed puncta similar to WT. One hypothesis is that FS mutants form overly stable complexes with WT DVL, impairing disassembly and signal transduction. This aligns with phenotypic differences observed in DVL null allele carriers, who lack the typical RS skeletal features (Carvalho et al. 2014; Zaveri et al. 2014), suggesting that FS variants may act via GOF mechanisms.

Given the dual roles of DVL in canonical and noncanonical WNT pathways, our findings support therapeutic strategies that aim to rebalance WNT signaling. For LOF alleles in WNT5A, ROR2, or FZD2, Wnt5a-derived hexapeptide Foxy-5 may restore PCP signaling (Safholm et al. 2008). For DVL-related RS, more nuanced interventions may be needed to modulate canonical activity—e.g., inhibitors targeting LRP or β-catenin interactions—or combined use with Foxy-5.

In summary, we provide evidence that the novel DVL C-terminus caused by frameshifting variants interferes with phosphorylation, subcellular localization, and WNT signaling activity conditioned upon WNT stimulus. Fig. 6 depicts a representative model for the potential molecular biological basis of this specific type of disease associated variant allele that generates a novel C-terminus of DVL, which affects the phosphorylation sites within the DVL C-terminus potentially leading to change in protein stability in response to WNT signaling. The mutant DVL with a novel C-termini may interact or competes with WT DVL, to form a hetero-supramolecular complex containing all three orthologs which likely interferes with the expected intracellular localization changes we observe under WNT stimulation. Moreover, co-expression of mutant alleles and WT alleles activate canonical WNT activity even in the absence of ligand stimulation.While *DVL1* and *DVL3* exhibited only modest activation,

**Fig. 6.**
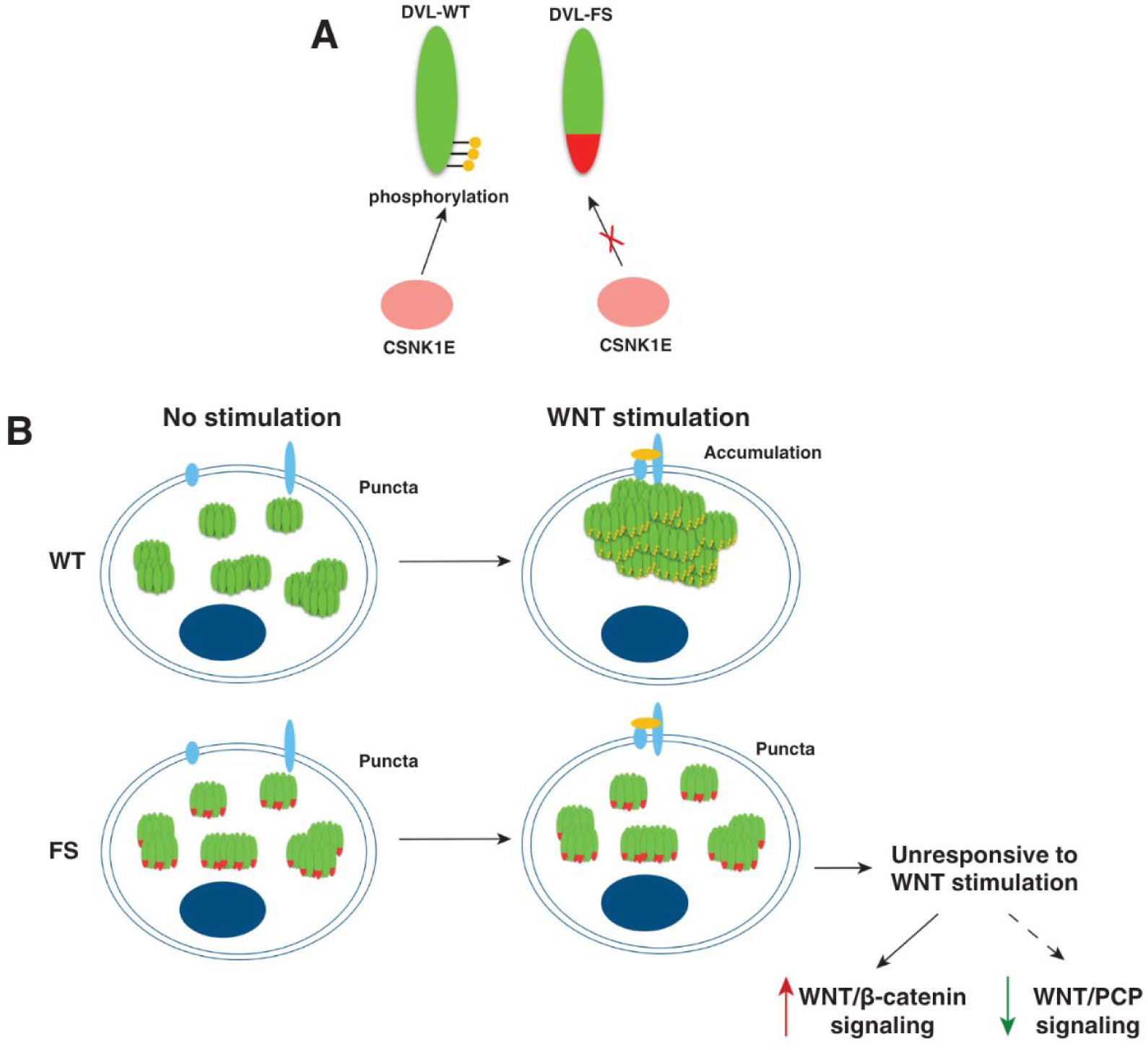
Novel C-terminal tails resulting from RS-frameshifting variants in *DVL* paralogs are not responsive to WNT stimulus. (A) WT DVL undergoes CSNK1E-mediated phosphorylation at its C-terminus, whereas DVL-FS lacks key residues at the C-termini which disrupts the expected mobility shift pattern. (B) Protein modeling predicts the replacement of > 82 aa at the C-termini of DVLs, charge distribution change and disruption of the IDR likely affecting their ability to form higher order structures and interactions with other proteins. *In vitro* experimental results here showed that DVLs with the replaced C-terminal: i. fail to exhibit the CSNK1E-induced phosphorylation-associated gel mobility shift observed in WT DVL, ii. do not alter cytoplasm localization under WNT ligand stimulus, and iii. display altered canonical WNT/β-catenin signaling dynamics and reduced responsiveness to WNT3A stimulation compared to WT DVL. Because DVL functions at the intersection of canonical and PCP WNT signaling pathways, the schematic incorporates a proposed reduction in PCP signaling (dashed arrow) based on established reciprocal regulation between canonical and PCP pathways. PCP pathway activity was not directly measured in this study, and therefore this component represents a model-based inference rather than an experimentally demonstrated effect. Green: WT DVL protein; Red: novel C-terminal tail caused by frameshifting variants observed in RS patient; Yellow dot: phosphorylation site at C-terminus; Orange ellipse: WNT ligand; Blue ellipse: WNT receptor and co-receptor; Dark blue ellipse: nucleus.

*DVL2* showed a more pronounced and statistically significant effect. This is consistent with the clinical observations of high bone density in individuals with *DVL1* frameshifting variants (Shayota et al. 2020). Therefore, the signaling perturbation we observe in vitro may reflect altered biological processes during development that contribute to bone malformations in DVL-related RS. This work may further our understanding of DVL’s role in the WNT pathway and human development, deepens our insights into underlying pathogenic disease mechanisms related to the WNT pathway, sheds light on -1 frameshifting alleles biological perturbation, and reveals novel genes function in skeletal formation and bone mass phenotype.

### Materials and Methods Cloning of plasmid constructs

As shown in Fig. 2, the N-terminal GFP tagged constructs containing human ‘wild-type’ (WT) DVL1 (DVL1-WT), frameshifting DVL1 (c.1519del, DVL1-FS), and the short version of DVL1 without C-terminal tail (c.1519insTAA, DVL1-ST) were obtained from Dr. Stephen P. Robertson’s lab (Bunn et al. 2015). The pCMV-HA mammalian expression vector containing an N-terminal hemagglutinin (HA) epitope tag was purchased from Clontech Inc. (Mountain View, CA, Cat. No. 631604). The N-terminal HA tagged constructs containing human WT DVL3 (DVL3-WT), mutant DVL3 (c.1749del, DVL3-FS2; c.1585del, DVL3-FS3) were generated by cloning using restriction endonucleases EcoRI and BglII. Constructs containing short version of DVL3 without C-terminal tail (c.1749insTAA, DVL3-ST2; c.1585insTAA, DVL3-ST3) were generated by using a Q5 site-directed mutagenesis kit following the manufacturer’s instructions (New England BioLabs Inc., Cat. No. E0552). The N-terminal Flag tagged constructs (vector: pcDNA3.1^+^/N-DYK) containing human ‘wild-type’ (WT) DVL2 (DVL2-WT) and +1 frameshifting mutant DVL2 (c.2105dup, DVL2-2105dup) were purchased from GenScript Inc. (Piscataway, NJ, Cat. No. SC1200, SC1017). Constructs containing -1 frameshifting mutant version of DVL2 (c.2105del, DVL2-2105del) and short version of DVL2 without C-terminal tail (c.2105insTAA, DVL2-ST) were generated by using a Q5 site-directed mutagenesis kit following the manufacturer’s instructions. The C-terminal Flag tagged construct containing human Casein kinase 1 Epsilon (CSNK1E-WT, vector: pcDNA3.1^+^/C-(K)DYK) was purchased from GenScript Inc. (Piscataway, NJ, Cat. No. OHu10985D). All constructs described in this study were verified by Sanger dideoxy capillary sequencing. All plasmids generated in this study, including WT, FS, and ST variants of DVL1/2/3, are available from the corresponding author upon reasonable request.

### Cell culture, transfection, and treatments

HEK293T cells were grown in Gibco Dulbecco’s Modified Eagle Medium (DMEM) with 4.5 g/L D-Glucose, L-Glutamine, 25mM HEPES, 10% Fetal bovine serum (FBS), 1X Antibiotic-Antimycotic. Cells were seeded in 24-well plates and were transfected the next day using Lipofectamine^TM^ 3000 reagent (ThermoFisher Scientific Inc., Waltham, MA, Cat. No. L3000015), following the manufacturer’s recommendations. Cells were incubated overnight after transfection, old medium was exchanged with new medium with or without recombinant Human/Mouse Wnt-5a or Recombinant Human Wnt-3a from Bio-techne Inc. (Minneapolis, MN. Cat. No. 645-WN-010, 5036-WN-010). Cells were harvested for immunoblotting or immunocytofluorescence 6-24 hours after medium change.

### Quantitative Reverse Transcription Polymerase Chain Reaction (qRT-PCR)

Total RNA was extracted from HEK293T cells using the RNeasy Midi kit (Qiagen, Cat. No. 74004) following the manufacturer’s protocol. Subsequently, cDNA was synthesized using 1 µg of RNA and the LunaScript RT SuperMix kit (New England Biolabs, Cat. No. E3010S). To quantify the expression of DVL 1, 2, and 3, a SYBR Green qPCR assay was performed using 1 µl of cDNA and the SYBR Green qPCR Master Mix (New England Biolabs, Cat. No. M3003L). The choice of *ACTB* (MIM 102630), encoding beta-actin, served as the endogenous control. The qPCR assay was conducted with the following cycling conditions: initial denaturation at 95°C for 60 seconds, followed by 40 cycles of denaturation at 95°C for 15 seconds, and extension for 30 seconds. The primers used are showing in Supplemental Table 1.

### Western blotting

Cells were transfected with corresponding constructs, incubated overnight, old medium was replaced with new medium. Cells were harvested 24 hours after medium change, washed with PBS, then centrifuged, pellets were collected. Samples were treated with RIPA or NP40 lysis buffer containing protease inhibitor cocktail (GenDepot Inc., Katy, TX, Cat. No. P3100), Halt Phosphatase Inhibitor (Thermo Scientific Inc., Rockford, IL, Cat. No.78420) shaken for 30 mins, then centrifuged 1000 xg at 4 C: for 10 mins. The supernatants were collected and stored at - 20 C:. Protein concentration was measured by using Pierce^TM^ BSA protein assay kit (Thermo Scientific Inc., Rockford, IL, Cat. No. NE173229) following the manufacturer’s instructions. Cell lysate samples were mixed with 4X Laemmli buffer (Bio-Rad Inc., Hercules, CA, Cat. No. 1610747) with 5% 2-Mercaptoethanol and were boiled for 5 mins at 95 C:. Samples were added to 7.5-10% TGX (Tris-Glycine eXtended) PreCast Stain-Free gel, run for ∼1.5 hour at 100V, wet transferred (60V, ∼1 hour) to PVDF membranes (Bio-Rad Inc., Hercules, CA, Cat. No. 162-0219). Membranes were incubated for 1 hour in blocking buffer (5% milk in TBST) or Tris Buffered Saline (TBS) with 5% (w/v) milk incubated overnight in primary antibodies at 4 C:. The next day membranes were washed 3 times for 15 mins with TBST, incubated in secondary antibodies for 1 hour at room temperature, washed 6 times for 15 mins with TBST. Membranes were incubated in SuperSignal^TM^ west pico PLUS chemiluminescent substrate (Thermo Scientific Inc., Rockford, IL, Cat. No. 34577) and scanned using FluorChem E system (Proteinsimple, San Jose, CA). Lambda Protein Phosphatase (New England Biolabs, Ipswich, MA, Cat. No. P0753S) used to check the phosphorylation state of CSK1e. The following antibodies were used: Anti-GFP (EMD Millipore Inc., Temecula, CA, Cat. No. AB10145, 1:5000 in bocking solution), Anti-HA (Cell signaling Inc., Danvers, MA, Cat. No. 3724, 1:1000 in TBST), DVL2 antibody (Cell Signaling Technology, Inc., Danvers, MA, Cat No. 3216, 1:1000 in blocking solution), Anti-GAPDH (Cell signaling Inc., Danvers, MA, Cat. No. 3683, 1:2000 in blocking solution), β-Actin (Cell signaling Inc., Danvers, MA, Cat. No. 6879, 1:1000 in blocking solution). All Western blots were performed in at least three independent biological replicates. Tag-specific antibodies (GFP, HA, and Flag) were used to distinguish individual DVL1, DVL2, and DVL3. The anti-DVL3 antibody recognizes all three endogenous DVL paralogs and was used in experiments that did not require paralog-specific discrimination.

### Immunocytofluorescence

Cells were seeded in 24-well plates with 12mm round cover glass, transfected with corresponding constructs, and incubated for 6-24 hours after medium change. Cells were rinsed with PBS, fixed in 4% paraformaldehyde in PBS for 20 mins, washed 3 times with PBST (PBS + 0.1% Triton X-100) for 5 mins, washed 2 times with PBSTB (PBST + 1% bovine serum albumin (BSA), Jackson ImmunoResearch Inc., West Grove, PA, Cat. No. 001-000-162) for 30 mins, blocked at least 30 mins in PBSTB + 5% Gibco^TM^ goat serum (ThermoFisher Scientific Inc., Waltham, MA, Cat. No. 16210-064), then incubated in PBSTB + primary antibody overnight at 4 C:.

The next day, cover glasses were washed with a few changes of PBSTB over the course of 1-2 hours, incubated with secondary antibody in PBSTB (1:500) for 1-2 hours, washed with a few changes of PBST over the course of 1-2 hours, then a final wash in PBST with DAPI (1:1000, Life technologies Inc., Eugene, OR, Cat. No. D1306) for 15 mins, incubated in PBS with Dylight^TM^ 650 Phalloidin (Cell signaling Inc., Danvers, MA, Cat. No.12956, 1:20), rinsed once with PBS, and mounted using SlowFade^TM^ Gold antifade reagent (Life technologies Inc., Eugene, OR, Cat. No. S36936) on glass slides, sealed with nail polish. The GFP-tagged constructs were imaged using live fluorescence signal rather than antibody-based detection. Cells were visualized using Zeiss 710 confocal microscope. Two independent experiments were performed for each condition, triplicate slides were generated for each condition per experiment. Five pictures were captured for each slide, positive cells per condition were analyzed and scored based on the pattern of DVL distribution. Confocal images were acquired with 40× and 63× objectives. All quantified localization analyses (puncta, even, accumulation) were performed under identical laser and detector settings. The following secondary antibodies was used: anti-Rb IgG (H+L) SuperClonalTM secondary antibody, alexa fluor 488 conjugate (Invitrogen Inc., Waltham, MA, Cat. No. 2059238), anti-Ms IgG (H+L) SuperClonalTM secondary antibody, alexa fluor 488 conjugate (Invitrogen Inc., Waltham, MA, Cat. No. A28175).

### TOPFlash Luciferase assays

Luciferase assays were performed on HEK293T cells. Transient transfections were performed at 60% confluence in 96-well plates (Corning Falcon Microplate, Cat. No. CLS353296). Cells were transfected using Lipofectamine 3000 (Invitrogen, Cat. No. L3000-008) with plasmids containing the wild type or variant *DVL* genes plus firefly reporter plasmids for SuperTopFlash (M50 Super 8x TOPFlash, Addgene plasmid #12456) and FOPFlash (M51 Super 8x FOPFlash, Addgene plasmid #12457) plasmid used as a negative control for “leaky” firefly luciferase transcription. A total of 100ng of plasmid was transfected in each well. WNT3A was used to measure WNT canonical activity and administered after 24 hours of cell plating. The Luciferase assays were performed after 48h of culture using Dual-luciferase reporter assay system (Promega, Cat. No. E1910). The Renilla plasmid is co-transfected in every well to normalize firefly luciferase activity. Biotek Synergy HT multi-mode microplate reader was used to read luminescence activity and GraphPad™ Prism™ software (La Jolla, CA) for analysis. Data represents average values of relative TOPFlash over FOPFlash ratios from three representative experiment performed in triplicate. Error bars indicate standard error of experiments performed in triplicate. Student t-test was used to determine statistical significance.

### *In silico* prediction of the effect of frameshift variants

The protein sequence alignment was performed using Cluster Omega Multiple Sequence Alignment (MSA, https://www.ebi.ac.uk/jdispatcher/msa/clustalo). The prediction of intrinsically disordered regions (IDRs) was conducted using IUPred2 (https://iupred2a.elte.hu/) (Meszaros et al. 2018). The amino acid sequence charges are prepared using EMBOSS charge tool (https://www.bioinformatics.nl/cgi-bin/emboss/charge) with a window size of 8. Further analysis was performed following the methods and code provided in previous work (Mensah et al. 2023).

Protein structure predictions were performed using AlphaFold2 Colab (Jumper et al. 2021).

### Statistical analysis

For cellular localization assays, data from two independent biological experiments were pooled, each containing three coverslips. Across replicates, more than 800 cells per condition were scored manually for three localization phenotypes (puncta, even, and accumulation). We analyzed puncta (or even) formation using mixed effect logistic regression models. We summarized the data as the number of punctate cells out of the total number of cells to evaluate whether treatments changed the probability of puncta (or even) formation while considering variability of technical replicates as the random effects. In the concentration (0-1 ng/uL) and treatment duration (0-24 hours) analyses we again employed mixed effect logistic regression. In the ligand–genotype analyses, we extended the binary mixed effects modeling to include additional explanatory factors including genotype (WT, FS, ST), ligand (none, WNT5A, WNT3A), and their interactions. *P*-values are computed by Wald tests. Post-hoc comparisons were performed against the appropriate reference condition and odds ratios with 95% confidence intervals are determined. For all post-hoc tests, *p*-values were corrected for multiple comparisons using the Dunnett method. For TOPFlash/FOPFlash luciferase assays, three independent biological replicates were performed, each measured in technical triplicate. Canonical WNT activity was quantified using the SuperTopFlash (STF) reporter, with FOPFlash serving as the negative control for nonspecific activation. STF/FOP ratios were used to correct for background luciferase activity. Relative STF/FOP values was analyzed using a linear mixed-effects model with log2-transformed activity as the outcome variable, genotype, ligand, and their interaction as fixed effects, and experiment as a random effect. *P*-values are computed by Wald tests. Post-hoc comparisons were performed against the appropriate reference condition and odds ratios with 95% confidence intervals are determined. For all post-hoc tests, p-values were corrected for multiple comparisons using the Dunnett method. For western blots, DVL proteins were analyzed, and band intensities were quantified using Fiji (ImageJ). For each DVL isoform, three independent biological replicates were performed. Relative band intensity was analyzed using a linear mixed-effects model, genotype, CSNK1E, and their interaction as fixed effects, and experiment as a random effect. Within each genotype, we performed pair-wise comparisons with *p*-values corrected by the Tukey-Kramer method. Statistical significance was defined as follows: *p* < 0.05 (\**), p* < 0.005 (**)*, p* < 0.0005 (***), and *p* < 0.0001 (****).

## Acknowledgments

The authors would like to thank Dr. Stephen P. Robertson who provided *DVL1* constructs that we used for the experiments, Dr. Yun Yan and Dr. Yikai Luo for helping review the imaging and statistical data.

## Statements & Declarations Funding

This work was supported in part by the US National Human Genome Research Institute (NHGRI)/ National Heart Lung and Blood Institute (NHLBI) grant number UM1HG006542 to the Baylor-Hopkins Center for Mendelian Genomics (BHCMG), the NHGRI Genome Research Elucidates Genetics of Rare disease (BCM-GREGoR) Program U01 HG011758, the National Institute of Neurological Disorders and Stroke (NINDS) R35 NS105078 (J.R.L.), the Eunice Kennedy Shriver National Institute of Child Health & Human Development (NICHD) R03 HD092569 (C.M.B.C and V.R.S.), and the National Institute of General Medical Sciences (NIGMS): R01 GM152556 (C.M.B.C). W-L.C. was supported by Cancer Prevention Research Institute of Texas (CPRIT) training Program RP140102. The work was supported in part by the Clinical Translational Core of Baylor College of Medicine Intellectual and Developmental Disabilities Research Center (P50HD103555) from the Eunice Kennedy Shriver NICHD. The content is solely the responsibility of the authors and does not necessarily represent the official views of the NIH.

## Competing Interest

J.A.G. have received travel support from Oxford Nanopore Technologies. Other authors declare no competing financial interests.

## Author contributions

Conceptualization was performed by Chaofan Zhang, Janson White, James R. Lupski and Claudia M.B. Carvalho, investigation and data analysis were performed by Chaofan Zhang, Rituparna Sinha Roy, Ming Yin Lun, Juliana F. Mazzeu, Janson White, Wu-Lin Charng, Nathaniel Peters, Jonas A. Gustafson, Harshini Iyer, Zain Dardas, James R. Lupski, and Claudia M.B. Carvalho, methodology and visualization were performed by Chaofan Zhang, Rituparna Sinha Roy, Ming Yin Lun, Chad A. Shaw, Hyun Kyoung Lee, V. Reid Sutton, James R. Lupski and Claudia M.B. Carvalho, supervision by Chad A. Shaw, Hyun Kyoung Lee, V. Reid Sutton, James R. Lupski and Claudia M.B. Carvalho. The first draft of the manuscript was written by Chaofan Zhang, and all authors commented on previous versions of the manuscript. All authors read and approved the final manuscript.

## Data Availability

The datasets generated during and/or analyzed during the current study are available from the corresponding author on reasonable request.

## Abbreviations

DEP: Dishevelled, Egl-10 and Pleckstrin domain
DRS: Dominant Robinow Syndrome
FS: frameshift
GFP: Green Fluorescent Protein
GOF: Gain-Of-Function
HPO: Human Phenotype Ontology
IDRs: intrinsically disordered regions
LP: Lambda phosphatase
MIM: Mendelian Inheritance in Man
PCP: Planar cell polarity
pLDDT: predicted Local Distance Difference Test
PDZ: acronym combining the first letters of three proteins — post synaptic density protein (PSD95), *Drosophila* disc large tumor suppressor (DlgA), and zonula occludens-1 protein (zo-1)
qRT-PCR: Quantitative Real-Time Reverse Transcription Polymerase Chain Reaction
RS: Robinow syndrome
ST: short
STF: SuperTopFlash
WT: Wild-Type

## Notes

### Summary of Updates

Added one author, Figure 3-6 updated, statistic analyses updated, supplemental files updated, manuscript details updated

