## Supplemental file for "Pathogenic *DVL* frameshifting variants in Robinow syndrome disrupt WNT signaling and cellular dynamics"

[1]Department of Molecular and Human Genetics, Baylor College of Medicine (BCM), Houston, TX 77030, USA; [2]Pacific Northwest Research Institute (PNRI), Seattle, WA 98122, USA; [3]University of Brasilia, Faculty of Medicine, Brasilia, 70910-900, Brazil; [4]Robinow Syndrome Foundation, Anoka, MN 55303, USA; [5]Instituto Nacional de Doenças Raras (InRaras), Universidade de Brasília, Brasília, Brazil; [6]Microscope Center, University of Washington, Seattle, WA 98195, USA; [7]Molecular and Cellular Biology Program, University of Washington, Seattle, Washington 98195, USA; [8]Department of Statistics, Rice University, Houston, Texas 77005, USA; [9]Department of Pediatrics, Section of Neurology, BCM, Houston, Texas 77030, USA; [10]Jan and Dan Duncan Neurological Research Institute, Texas Children's Hospital, Houston, Texas 77030, USA; [11]Department of Neuroscience, BCM, Houston, Texas 77030, USA; [12]Texas Children's Hospital, Houston, TX 77030, USA; [13]Human Genome Sequencing Center, BCM, Houston, TX 77030, USA; [14]Department of Pediatrics, BCM, Houston, TX 77030, USA; [15]These authors contribute equally to this work.

*Corresponding author:

Claudia M. B. Carvalho, PhD

Associate Investigator

Pacific Northwest Research Institute (PNRI)

720 Broadway, Seattle, WA, 98122

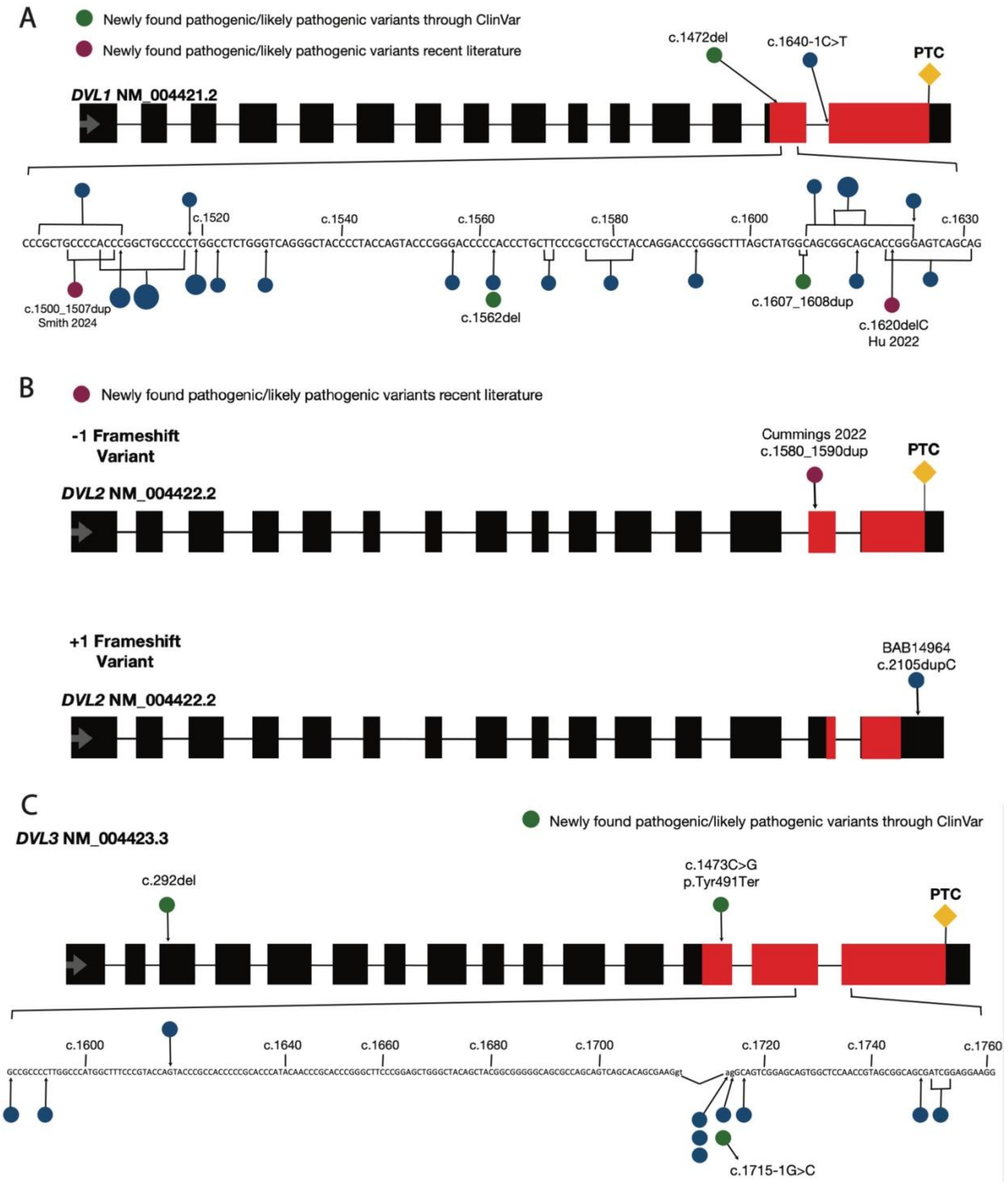
A
Newly found pathogenic/likely pathogenic variants through ClinVar
Newly found pathogenic/likely pathogenic variants recent literature
c.1472del
c.1640-1C>T
PTC
DVL1 NM_004421.2
c.1520
c.1540
c.1560
c.1580
c.1600
c.1630
c.1500_1507dup
Smith 2024
c.1562del
c.1607_1608dup
c.1620delC
Hu 2022
B
Newly found pathogenic/likely pathogenic variants recent literature
-1 Frameshift Variant
Cummings 2022
c.1580_1590dup
PTC
DVL2 NM_004422.2
+1 Frameshift Variant
BAB14964
c.2105dupC
DVL2 NM_004422.2
C
Newly found pathogenic/likely pathogenic variants through ClinVar
DVL3 NM_004423.3
c.292del
c.1473C>G
p.Tyr491Ter
PTC
c.1600
c.1640
c.1660
c.1680
c.1700
c.1720
c.1740
c.1760
c.1715-1G>C

**Supplemental Figure 1. Map location of pathogenic/likely pathogenic DNA variants affecting *DVL1*, *DVL2* and *DVL3* in individuals diagnosed with DRS.** Chromosome and cytogenetic interval locations of the canonical transcripts are provided, individual exons (black rectangles) on a horizontal line are drawn to represent gene structure. Previously described variants are represented by blue circles, newly variants are represented by green circles (ClinVar) or by red circles (literature) (1-3); the size of the circles is proportional to the number of recurrent variants in unrelated individuals. **A. Variants within the coding region of *DVL1*:** Variants consist of mostly of small insertions or deletions (indels), except for one splicing variant. All variants are predicted to lead to -1 frameshifting of the reading frame. Zoomed-in view shows the bases affected by variants in exon 14 (i.e., penultimate exon). A premature termination codon (PTC) created by -1 frameshifting variants is displayed by an orange diamond. Location of the predicted starting amino acid corresponding to the minimum NMD-escape region is at 1466 aa in this transcript. **B. Variant within the coding region of *DVL2*:** Only two unrelated individuals with DRS have been reported carrying likely pathogenic variants, both are indels. One individual carries a *de novo* 1 bp duplication at the position c.2105dupC predicted to lead to a +1 frameshifting and a longer protein (4), whereas the second variant consisting of a 11 bp duplication at position c.1580_1590dup, is predicted to lead to -1 frameshifting with a PTC at exon 15 similar to *DVL1* variants. The location of predicted starting amino acid of the NMD-escape region is 1546 to 2140 aa for -1 frameshifting variants, whereas 1708 to 1967 aa for +1 frameshifting variants. **C. Variants within the coding region of *DVL3*:** Most pathogenic/likely pathogenic variants reported in *DVL3* consist of indels, except for five splicing variants and a nonsense variant. Analogous to *DVL1*, all pathogenic/likely pathogenic variants are predicted to lead to -1 frameshifting of the reading frame, except the nonsense variant which is predicted to lead to undergo degradation by the NMD machinery. PTCs created by -1 frameshifting variants are displayed by an orange diamond. Zoomed-in view shows the bases affected by variants in exon 14 and 15. The location of predicted starting amino acid of the NMD-escape region is 1416 aa.

**A**

```
DVL1_WT            DTLAPLPHPAAPWPLGQGYPYQYPGPPPCFPPAYQDPGFSYGSGSTGSQQSEGSKSSGSTRSSRRAPGREKERRAAGAGGSGSESDHT

c.1619_1631del*    DTLAPLPHPAAPWPLGQGYPYQYPGPPP--CFPPAYQDP--GFSYGSGSRVKGAKAVGPPGAAAGPRAVRRSVGRRELGAVAVNRITR
c.1519del          DTLAPLPHPAAPGLWVRATPTSTRDPHPASRLPTRTRALAMAAAAPGVSRVKGAKAVGPPGAAAGPRAVRRSVGRRELGAVAVNRITR
c.1496_1508del#    DTLAR----LPPGLWVRATPTSTRDPHPASRLPTRTRALAMAAAAPGVSRVKGAKAVGPPGAAAGPRAVRRSVGRRELGAVAVNRITR
                   ****        *     :. * .  .* *   :*    :      . :   . ****************************************

DVL1_WT            APSGVGSSWRERPAGQLSRGSSPRSQASATAPGLPPPHPTTKAYTVVGGPPGGPPVRELAAVPPELTGSRQSFQKAMGNPCEFFVDIM

c.1619_1631del*    HRVGWGAAGESVRPASSAVAAAHAVRPRLPPRGSPRPTPRPRPIQWWGGHPGDPLSGSWLPSPRN*
c.1519del          HRVGWGAAGESVRPASSAVAAAHAVRPRLPPRGSPRPTPRPRPIQWWGGHPGDPLSGSWLPSPRN*
c.1496_1508del#    HRVGWGAAGESVRPASSAVAAAHAVRPRLPPRGSPRPTPRPRPIQWWGGHPGDPLSGSWLPSPRN*
                   *****************************************************************
```

**B**

```
DVL2_WT          PPGAPPVRDLGSVPPELTASRQSFHMAMGNPSEFFVDVM
DVL2_c.2105del   PPGALQSETWALCPQN
DVL2_c.2105dup   PPGAPSSQRPGLCAPRTDSQPPKLPHGHGQSQRVLCGCYVAHCGARLGRALLVCDWVSWPSCACSYSAWAQPTSCCHTGDCGHCDSHQQCLVPPPSLRGRQGTFDYF
```

**C**

```
DVL3_WT                LPHPGAAPWPMAFPYQYPPPPHPYNPHPGFPELGYSYGGGSASSQHSEGSRSSGSNRSGSDRRKEKDPKAGDSKSGGSGSESDHTTRSSLRGPRERA

DVL3_c.1749del         LPHPGA--APWPMAFPYQYPPPPHPYNPHPGFPELGYSYG----GGSASSQHSEGSRSSGSNRSGRIGGRRRTRRPGTPSPGAAAANRTTPHAAACG
DVL3_c.1751_1754del#   LPHPGA--APWPMAFPYQYPPPPHPYNPHPGFPELGYSYG----GGSASSQHSEGSRSSGSNRSGS-GGRRRTRRPGTPSPGAAAANRTTPHAAACG
DVL3_c.1585del*        LPHPGPPLGPWLSRTSTRHPRT-H-----TTRTRASRSWATATAGAAPAVSTAKAVGAVAPTVAAAIGGRRRTRRPGTPSPGAAAANRTTPHAAACG
DVL3_c.1617del         LPHPGAAPWPMAFPYHTRHPRT-H-----TTRTRASRSWATATAGAAPAVSTAKAVGAVAPTVAAAIGGRRRTRRPGTPSPGAAAANRTTPHAAACG
                       *****    *       ::*   *          . . *:.    *.: : . ::.  : . . :.  ******************************

DVL3_WT                PSERSGPAASEHSHRSHHSLASSLRSHHTHPSYGPPGVPPLYGPPMLMMPPPPAAMGPPGAPPGRDLASVPPELTASRQSFRMAMGNPSEFFVDVM

DVL3_c.1749del         GRGSGRPASAQGRRPASTATAATIPWPAAFAATTHTRATVLPECPLSTAPPC
DVL3_c.1751_1754del#   GRGSGRPASAQGRRPASTATAATIPWPAAFAATTHTRATVLPECPLSTAPPC
DVL3_c.1585del*        GRGSGRPASAQGRRPASTATAATIPWPAAFAATTHTRATVLPECPLSTAPPC
DVL3_c.1617del         GRGSGRPASAQGRRPASTATAATIPWPAAFAATTHTRATVLPECPLSTAPPC
                       ****************************************************
```

**D**

```
DVL1_WT  LPHP-AAPWPLGQGYPYQYPGPPPCF----PPAYQDPGFSYGSGSTGSQQSEGSKSSGSTRSSRRAPG--REKER-RAAGAGGSGSESDHTAPSGVGSSWRERP
DVL2_WT  APLPGATPWPLLPTFSYQYP-APHPYSPQPPPYHELSSYTYGGGSASSQHSEGSRSSGSTRSDGGAGRTGRPEERAPESK-SGSGSESEPSSRGGSLRRGGEAS
DVL3_WT  LPHPGAAPWPMAF--PYQYPPPPHPYNPHP--GFPELGYSYGGGSASSQHSEGSRSSGSNRSGSDRRKEKDPK--AGDSKSGGSGSESDHTTRS-SLRGPRERA
          * * *:***:     ***** *  :        .   .::**.**:.**:****:****.**,          :     :  .******: :: .       *

DVL1_WT  AGQLSRGSSP----------------RSQA----SATAPGLPPPH---------PTTKAYTVVGGPPGGPPVRELAAVPPELTGSRQSFQKAMGNPCEFFVDIM
DVL2_WT  G--TSDGGPPPS----RGSTGGAPNLRAHPGLHPYGPPPGMALPYNPMMVVMMPPPPPPVPPAVQPPGAPPVRDLGSVPPELTASRQSFHMAMGNPSEFFVDVM
DVL3_WT  P--SERSGPAASEHSHRSHHSLASSLRSHHTHPSYG-PPGVPPLYGPPMLMMPPP-----PAAMGPPGAPPGRDLASVPPELTASRQSFRMAMGNPSEFFVDVM
            . ..                    *::      .  **     :          *        .  ***.** *:*.:******.*****: *****.*****:*
```

**E**

```
DVL1_1519del  LPHPAAP-GLWVRATPTSTRDPHPASRLPTRTRALAMAAAAPGVSRVKGAKAVGPPGA-AAGPRAVRRSVGRRELGAVAVNR
DVL3_1585del  LPHPGPPLGPWLSRTSTRHPRTHTTRTRASRSWATATAGAAPAVSTAKAVGAVAPTVAAAIGGRRRTRRPGTPSPGAAAANR
              ****. * * *:  * *     * :    :*: * * *.***.** .*.. **.*  * * * *   *  *  . **.*.**

DVL1_1519del  ITRHRVGWGAAGES-------VRPASSAVAAAHAVRPRLPPRGSPRPTPRPRPIQWWGGHPGDPLS-GSWLPS-PRN-----
DVL3_1585del  TTPHAAACGGRGSGRPASAQGRRPASTATA---------------------ATIPWPAAFAATTHTRATVLPECPLSTAPPC
               * * .. *. *..        ****:*.*                     * * ... .    : .: **. *.
```

Red: Small and hydrophobic
Blue: Acidic
Magenta: Basic
Green: Hydroxyl, sulfhydryl, and amine residues

- "*": Identical conserved residues
- ":": Strongly similar residues
- ".": Weakly similar properties
- No symbol: Residues that are not conserved

**F**

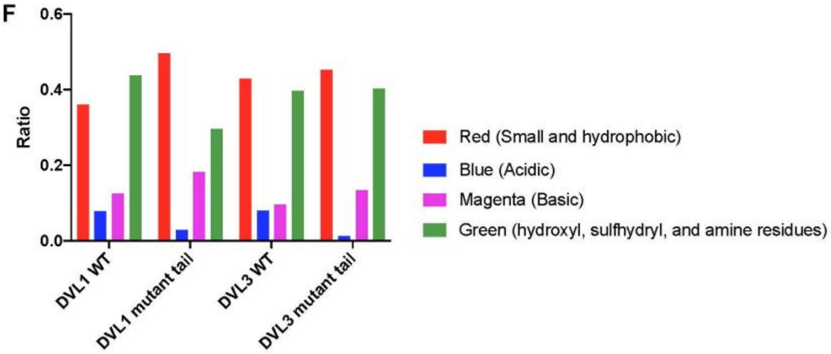

**Supplemental Figure 2. Amino acids residues comparison between WT and mutant DVL C-termini regarding chemical properties and conservation patterns.** WT DVL C-termini are compared to DVL variants investigated in this study, both of which compared to *DVL1* and *DVL3* variants from Supplemental Figure 1 predicted to result in the longest (indicated by a star) and shortest (indicated by a hash mark) C-terminal tails. **A.** C-terminal region of WT DVL1 consisting of 171 amino acids is compared to the DVL1 variant C-terminal (c.1619_1631del, c.1519del, c.1496_1508del) involving 144 to 148 amino acids. All pathogenic/likely pathogenic frameshifting variants in *DVL1* (NM_004421.2) are predicted to generate the same mutant C-terminal tail containing a minimum of 106 amino acids replacing the WT residues and resulting in a loss of 23 amino acids compared to WT DVL1. **B.** C-terminal region of WT DVL2 consisting of 39 amino acids is compared to the predicted +1 frameshift variant (107 amino acids) and -1 frameshift variant (16 amino acids) of *DVL2* (NM_004422.2). The +1 frameshift produces a protein extended by 68 amino acids with a novel mutant tail, whereas the -1 frameshift results in a protein shortened by 23 amino acids. **C.** The C-terminal region of WT DVL3 consisting of 193 amino acids is compared to the variant DVL3 (c.1749del, c.1751_1754del, c.1585del, c.1617del) involving 142 amino acids. All pathogenic/likely pathogenic frameshifting variants in *DVL3* (NM_004423.3) are predicted to produce the same mutant C-terminal tail containing a minimum of 82 amino acids replacing the WT residues and resulting in a loss of 44 amino acids compared to the WT DVL3. **D.** Amino acid residue conservation comparison of the C-termini of WT DVL1, DVL2, and DVL3. **E.** Amino acid residue conservation comparison of selected mutant forms of DVL1 and DVL3. **F.** Bar plot representing the chemical ratio alteration of the amino acid residues in the C-termini of WT and mutant DVL1 (c.1519del, 143 amino acids) and DVL3 (c.1585del, 142 amino acids).

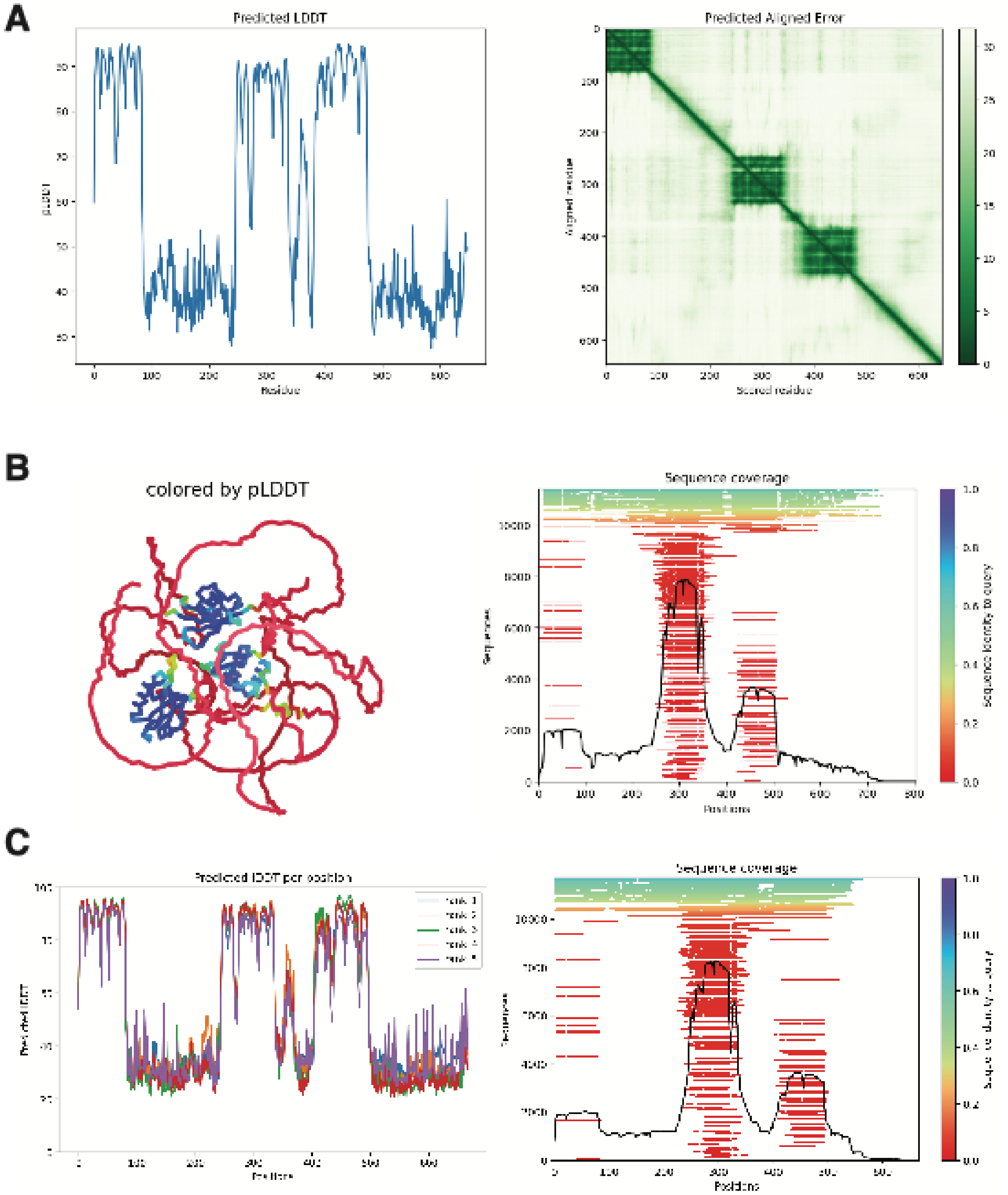


**Supplemental Figure 3. AlphaFold2 quality assessment of RS-associated *DVL* frameshift variants. A**. *DVL1* c.1519del, **B**. *DVL2* c.2105dup, and **C**. *DVL3* c.617del models are shown. Predicted local distance difference test (pLDDT) scores across the protein sequence demonstrate high confidence within structured domains and markedly reduced confidence in the altered C-terminal regions. Predicted aligned error (PAE) and sequence coverage plots further indicate limited structural confidence and lack of homologous template support in the frameshifted C-termini. These results highlight the intrinsic limitations of structure prediction for the abnormal C-terminal tails, which lack homology and are predicted to be highly disordered.

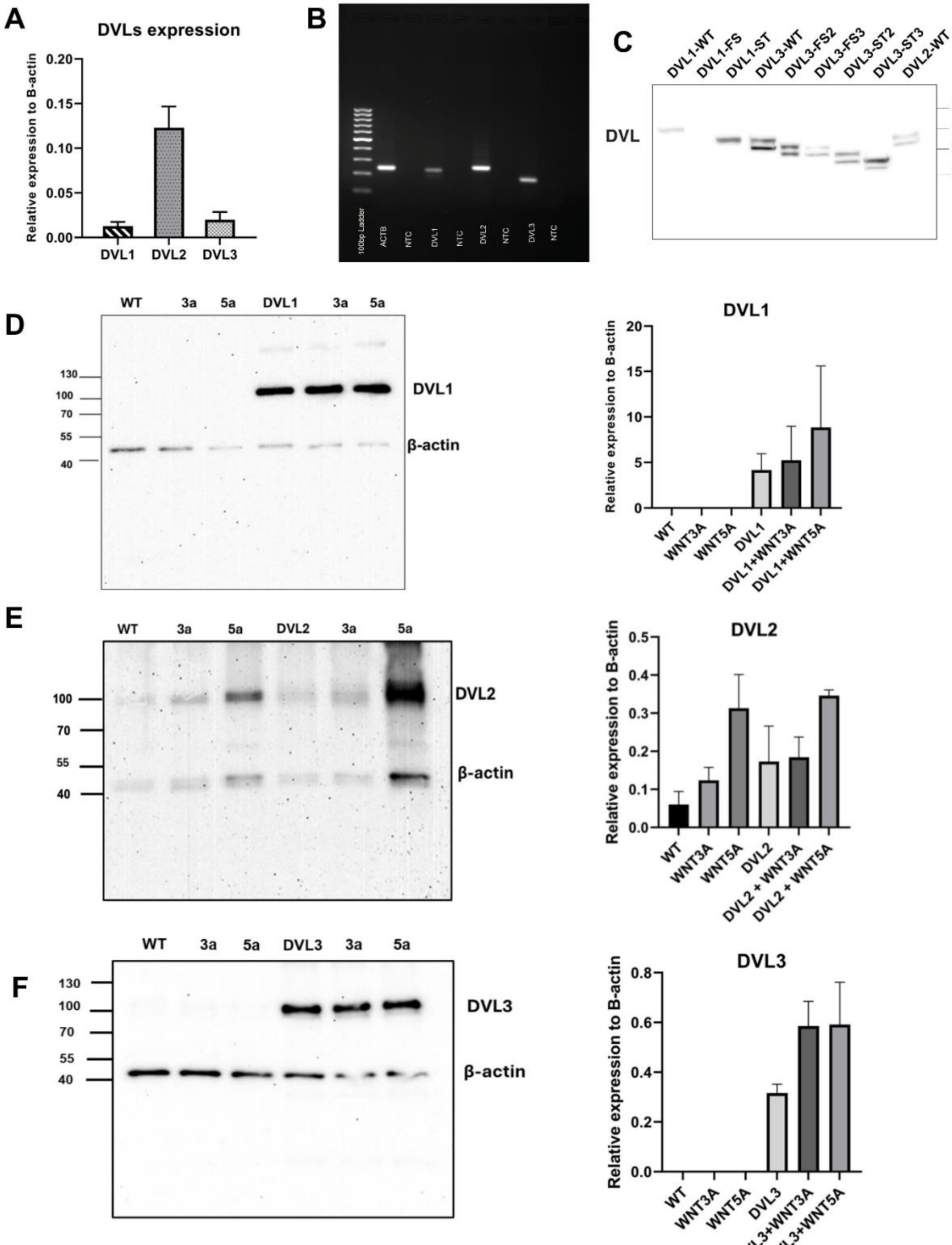

A
DVLs expression
Relative expression to B-actin
0.20
0.15
0.10
0.05
0.00
DVL1
DVL2
DVL3
B
100bp Ladder
ACTB
NTC
DVL1
NTC
DVL2
NTC
DVL3
NTC
C
DVL1-WT
DVL1-FS
DVL1-ST
DVL3-WT
DVL3-FS2
DVL3-FS3
DVL3-ST2
DVL3-ST3
DVL2-WT
DVL
150kD
100kD
75kD
50kD
D
WT
3a
5a
DVL1
3a
5a
130
100
70
55
40
DVL1
β-actin
DVL1
Relative expression to B-actin
20
15
10
5
0
WT
WNT3A
WNT5A
DVL1
DVL1+WNT3A
DVL1+WNT5A
E
WT
3a
5a
DVL2
3a
5a
100
70
55
40
DVL2
β-actin
DVL2
Relative expression to B-actin
0.5
0.4
0.3
0.2
0.1
0.0
WT
WNT3A
WNT5A
DVL2
DVL2 + WNT3A
DVL2 + WNT5A
F
WT
3a
5a
DVL3
3a
5a
130
100
70
55
40
DVL3
β-actin
DVL3
Relative expression to B-actin
0.8
0.6
0.4
0.2
0.0
WT
WNT3A
WNT5A
DVL3
DVL3+WNT3A
DVL3+WNT5A

**Supplemental Figure 4. Investigation of endogenous expression of DVL RNA and proteins in HEK293T cells and overexpression system.** Quantitative Reverse Transcription Polymerase Chain Reaction (qRT-PCR) was performed to assess the mRNA expression of *DVL1*, *2*, and *3* in relation to the reference gene *ACTB* (beta-actin) (**A, B**). **A.** Comparative analysis of *DVL* gene expression normalized to *ACTB*. *DVL1*, *DVL2*, and *DVL3* exhibited expression levels approximately 1.28%, 12.29%, and 1.99% of *ACTB*, respectively. **B.** Agarose gel (1%) electrophoresis showing single amplification products for each primer set confirming the specificity of the primers to each *DVLs*. Lane 1: GeneRuler 100 bp DNA ladder (ThermoScientific), lane 2: *ACTB*, lane 3: no template control for *ACTB*, lane4: *DVL*1, lane 5: no template control for *DVL*1, lane 6: *DVL*2, lane 7: no template control for *DVL*2, lane 8: *DVL3*, lane 9: no template control for *DVL3*. **C.** Western blot analysis of HEK 293T cells overexpressing different *DVL* constructs. The anti-DVL3 antibody (Thermo Scientific Inc., Rockford, IL, Cat. No. PA5-27723) was utilized, capable of detecting the wild-type versions of DVL1, DVL2, and DVL3 proteins, allowing for detection of endogenous DVL proteins. Western blot analysis of total protein extracted from HEK293T cells, either untransfected or transfected with GFP-tagged DVL1-WT (**D**), Flag-tagged DVL2-WT (**E**), or HA-tagged DVL3-WT (**F**), with or without treatment using WNT3A or WNT5A recombinant proteins. The relative expression levels of DVL1 protein were normalized to beta-actin and are presented as a bar graph. No statistical significance was reached (p-value threshold set at 0.05).

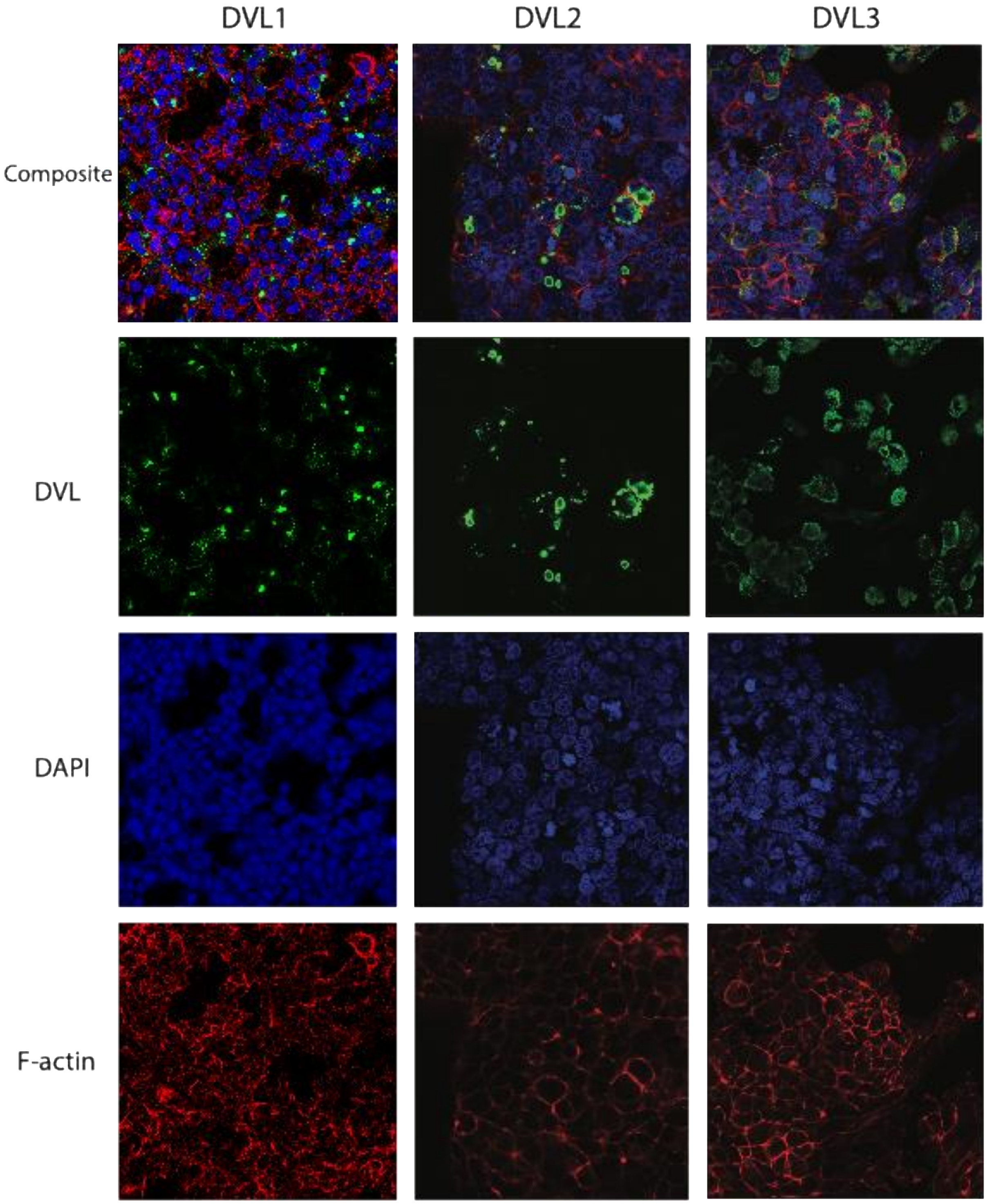
DVL1
DVL2
DVL3
Composite
DVL
DAPI
F-actin

**Supplemental Figure 5. Full-field images of immunocytofluorescence experiment for cells overexpressing WT GFP-tagged DVL1, FLAG-tagged DVL2, and HA-tagged DVL3 proteins.** The secondary anti-Rb IgG (H+L) SuperClonalTM secondary antibody, alexa fluor 488 conjugate antibody was used. Green: WT DVL1, DVL2, DVL3; Red: 650 Phalloidin, label of cytoskeleton; Blue: DAPI, label of nucleus. Confocal images were acquired with and 63× objectives.

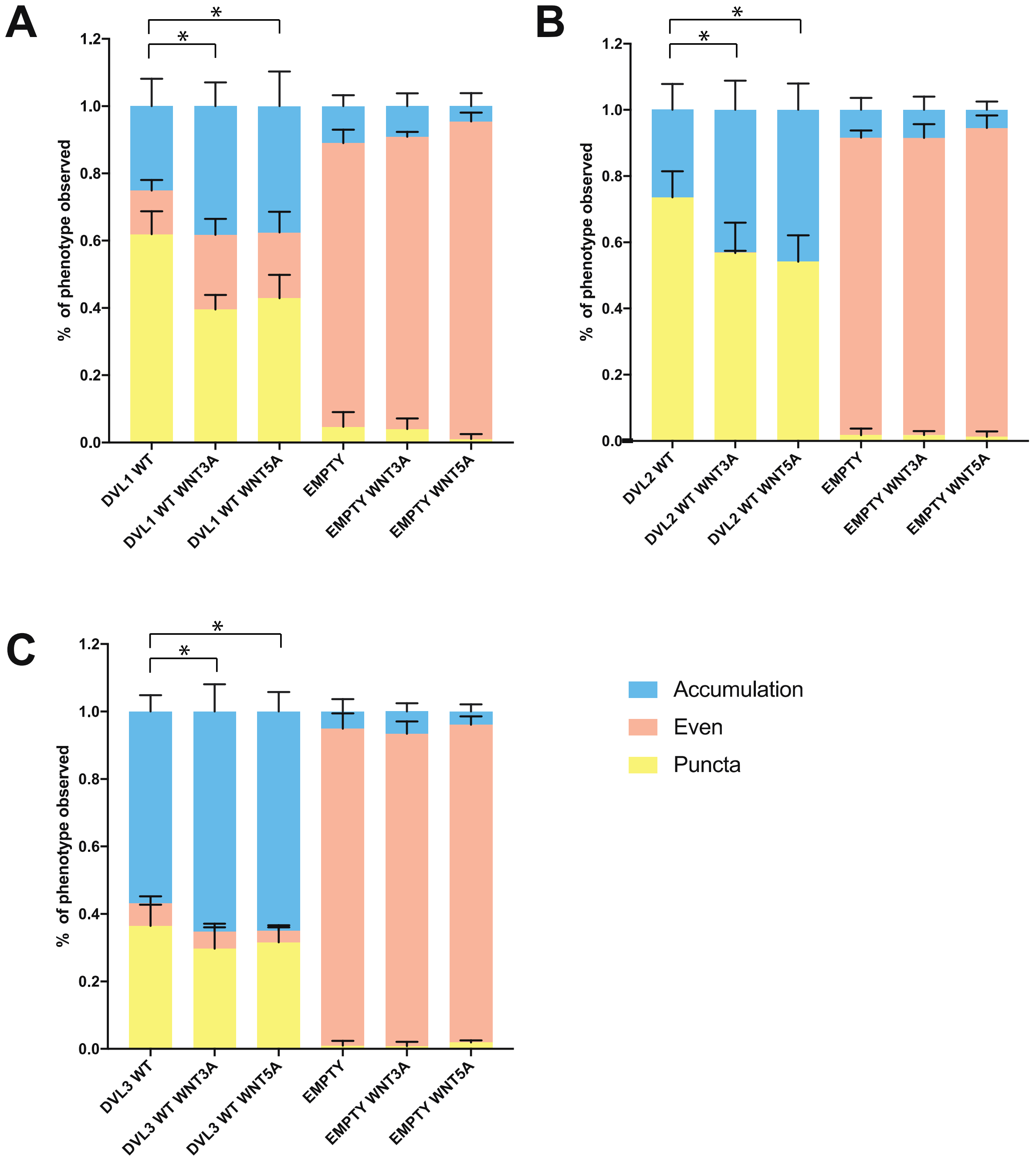


**Supplemental Figure 6. Subcellular localization of DVL and empty vectors**

**A.** HEK293T cells were transfected with GFP-tagged *DVL1* WT construct or GFP tagged empty construct, with or without treatment of 0.2 ng/uL of WNT3A or WNT5A recombinant protein, incubated for 6 hours. The ratio of distribution patterns of DVL1-GFP or GFP was calculated and analyzed. **B.** HEK293T cells were transfected with Flag-tagged *DVL2* WT construct or Flag tagged empty construct, with or without treatment of 0.2 ng/uL of WNT3A or WNT5A recombinant protein, incubated for 6 hours. The ratio of distribution patterns of DVL2-Flag or Flag was calculated and analyzed. **C.** HEK293T cells were transfected with HA-tagged *DVL3* WT construct or HA tagged empty construct, with or without treatment of 0.2 ng/uL of WNT3A or WNT5A recombinant protein, incubated for 6 hours. The ratio of distribution patterns of DVL3-HA or HA was calculated and analyzed.

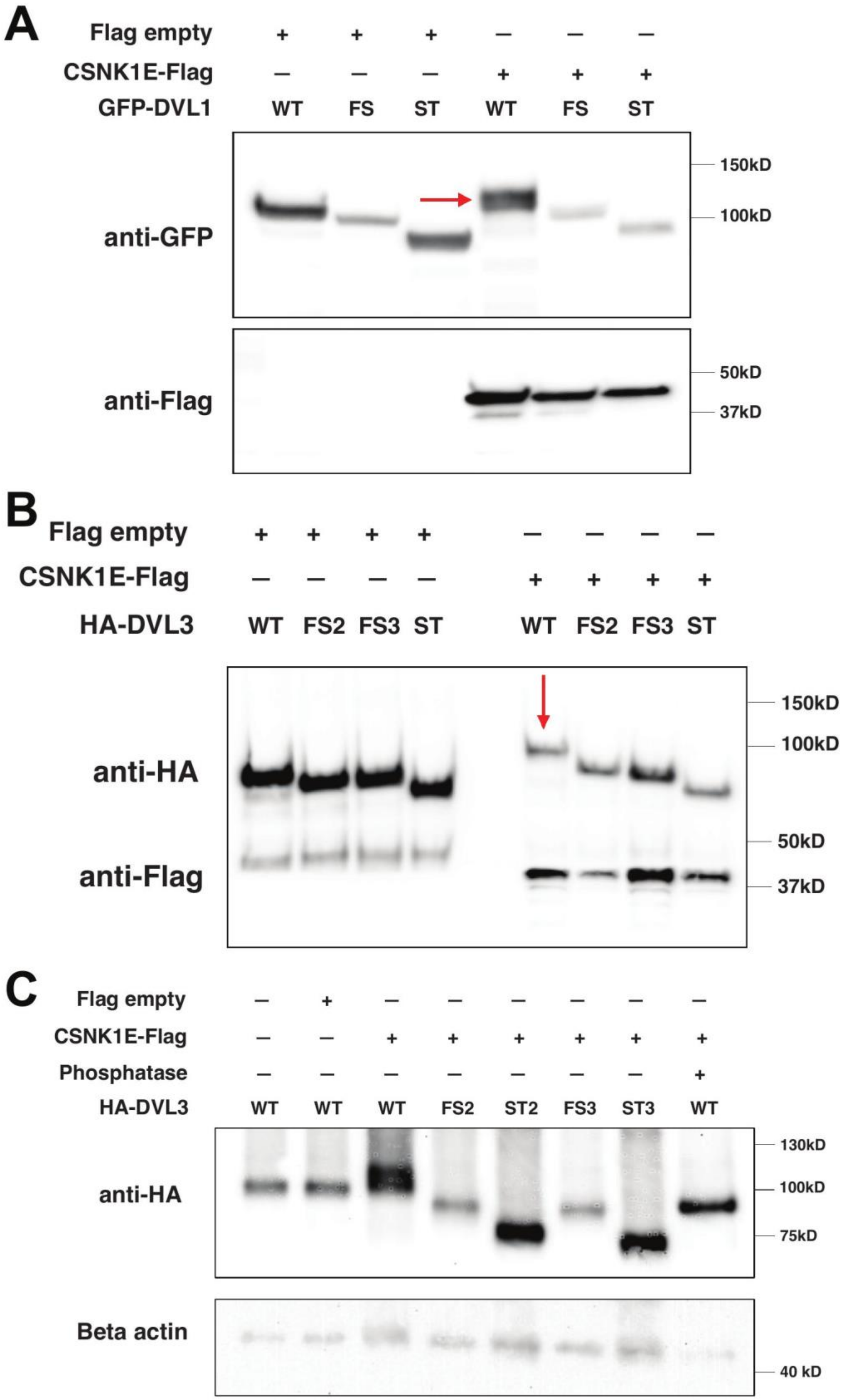
A
Flag empty
CSNK1E-Flag
GFP-DVL1
WT FS ST WT FS ST
anti-GFP
anti-Flag
150kD
100kD
50kD
37kD
B
Flag empty
CSNK1E-Flag
HA-DVL3
WT FS2 FS3 ST WT FS2 FS3 ST
anti-HA
anti-Flag
150kD
100kD
50kD
37kD
C
Flag empty
CSNK1E-Flag
Phosphatase
HA-DVL3
WT WT WT FS2 ST2 FS3 ST3 WT
anti-HA
Beta actin
130kD
100kD
75kD
40 kD

**Supplemental Figure 7: Impact of DVL constructs on CSNK1E induced electrophoretic migration and phosphorylation-dependent shift of DVL1 and DVL3.**

**A.** Western blot result for whole proteins extracted from HEK293T cells transfected with GFP-tagged DVL1-WT, DVL1-FS, DVL1-ST alone, or co-transfected with Flag-tagged CSNK1E or Flag empty construct with the DVL1 constructs. **B, C.** Western blot results for whole proteins extracted from HEK293T cells transfected with HA-tagged DVL3-WT, DVL3-FS2, DVL3-FS3, DVL3-ST alone, or co-transfected with Flag-tagged CSNK1E or Flag empty construct with the DVL3 constructs. Lambda phosphatase was added to confirm the migration is caused by phosphorylation.

Supplemental Table 1. Primers used for qRT-PCR

| Genes | Primers |
|---|---|
| *DVL1* | Forward 5’-AGACGGCGGCATCTACATTG-3’ |
| | Reverse 5’-GACGGTGAAGTAGCTTCGGG-3’ |
| *DVL2* | Forward 5’-GGAGAGTACCAGCCTGGGG-3’ |
| | Reverse 5’-GCTCTGGCCAACAATGGAGAT-3’ |
| *DVL3* | Forward 5’-CTTCCCTGCATACGGCATGA-3’ |
| | Reverse 5’-CGGACCTCCAACCCTGATTC-3’ |
| *ACTB* | Forward 5’-CCTGAAGTACCCCATCGAGC-3’ |
| | Reverse 5’-AGAGGCGTACAGGGATAGCA -3’ |
